# Choroid Plexus Modulates Subventricular Zone Adult Neurogenesis and Olfaction Through Secretion of Small Extracellular Vesicles

**DOI:** 10.1101/2023.03.16.532966

**Authors:** Luke L. Liu, Jonathan Shannahan, Wei Zheng

## Abstract

The choroid plexus (CP) in brain ventricles secrete cerebrospinal fluid (CSF) that bathes the adjacent subventricular zone (SVZ); the latter is the largest neurogenic region in adult brain harboring neural stem/progenitor cells (NSPCs) and supplies newborn neurons to the olfactory bulb (OB) for normal olfaction. We discovered the presence of a CP-SVZ regulatory (CSR) axis in which the CP, by secreting small extracellular vesicles (sEVs), regulated adult neurogenesis in the SVZ and maintained olfaction. The proposed CSR axis was supported by 1) differential neurogenesis outcomes in the OB when animals treated with intracerebroventricular (ICV) infusion of sEVs collected from the CP of normal or manganese (Mn)-poisoned mice, 2) progressively diminished SVZ adult neurogenesis in mice following CP-targeted knockdown of SMPD3 to suppress CP sEV secretion, and 3) compromised olfactory performance in these CP-SMPD3-knockdown mice. Collectively, our findings demonstrate the biological and physiological presence of this sEV-dependent CSR axis in adult brains.

**Highlights:**

1. CP-secreted sEVs regulate adult neurogenesis in the SVZ.
2. CP-secreted sEVs modulate newborn neurons in the OB.
3. Suppression of sEV secretion from the CP deteriorates olfactory performance.

## 1. Introduction

The choroid plexus (CP) produces and secretes cerebrospinal fluid (CSF) which transports nutrients, growth factors and regulatory elements to the central milieu to nourish brain cells and ensure normal brain functions (Fultz et al. 2019; Lehtinen et al. 2011; Zheng and Chodobski 2005). Immediately adjacent to the CP is the subventricular zone (SVZ), the largest neurogenic niche in the adult vertebrate brain, located in the walls of lateral ventricles. The SVZ harbors neural stem/progenitor cells (NSPCs) and supplies newborn neurons to the olfactory bulb (OB) for normal olfaction (Lim and Alvarez-Buylla 2016). The interaction between the CP and SVZ has recently attracted attention due to the CP’s regulation of NSPC dynamics in the adult brain (Lepko et al. 2019; Liu et al. 2022b; Planques et al. 2019; Silva-Vargas et al. 2016) as well as in the infant brain (Lehtinen et al., 2011). The concept of a **C**horoid plexus-**S**ubventricular zone **R**egulatory (CSR) axis, in which the regulatory signals delivered from the CP to the SVZ mediate NSPC proliferation, migration and differentiation in the SVZ-rostral migratory stream (RMS)-OB system, remain elusive. Additionally, less is known regarding what moiety secreted by the CP into the CSF may transmit the regulatory signals targeted towards the SVZ. Hence, thorough understanding of this axis helps address the olfactory disorder observed in numerous neurological conditions and facilitate therapeutic interventions.

Early studies from this lab along with cumulative data in literature have established the CP, which receives the most abundant blood flow within brain, serves as the target for insults from the circulation, such as peripheral inflammation (Balusu et al. 2016; Carloni et al. 2021), viral infection (Schwerk et al. 2015; Yang et al. 2021), and neurotoxicant exposure (Ingersoll 1995; Zheng 2001; Zheng et al. 1991). Specifically, exposure to toxic metal manganese (Mn) is known to cause Parkinsonian disorders among Mn-exposed welders and smelters (O’Neal and Zheng 2015), lead to extensive Mn accumulation in the CP and CSF (Zheng et al. 2009), cause aberrant adult neurogenesis in the SVZ (Fu et al. 2015, 2016), and induce olfactory dysfunction (Werner and Nies 2018). Thus, it seemed highly likely that the CP, in response to environmental insult in the systemic circulation, may mediate SVZ activities via this proposed CSR axis, leading to early signs and symptoms of Mn-associated neurodegeneration.

Small extracellular vesicles (sEVs, diameter < 200 nm) secreted by host cells carry distinct cellular cargos mediating intercellular communication (Corrado et al. 2013; Gurung et al. 2021; Rajput et al. 2022). The CP exhibits a robust expression of sEV-related markers (Dani et al. 2021; Lein et al. 2007). Considering the proximity between the CP and SVZ, the sensitivity of SVZ NSPCs to CSF signals, and abundant sEVs in the CP, we proposed the CP produces and secretes sEVs into the CSF; the SVZ senses these signals carried by the sEVs and facilitates neurogenesis in the SVZ-RMS-OB system. Overall, the data presented in this study through molecular, toxicological, and neurobehavioral characterizations support a physiologically viable CP-SVZ regulatory (CSR) axis mediated by sEVs.

## 2. METHODS AND MATERIALS

### 2.1. Chemicals and reagents

Chemicals and reagents used in this study were listed in Table S1.

**Table S1.**
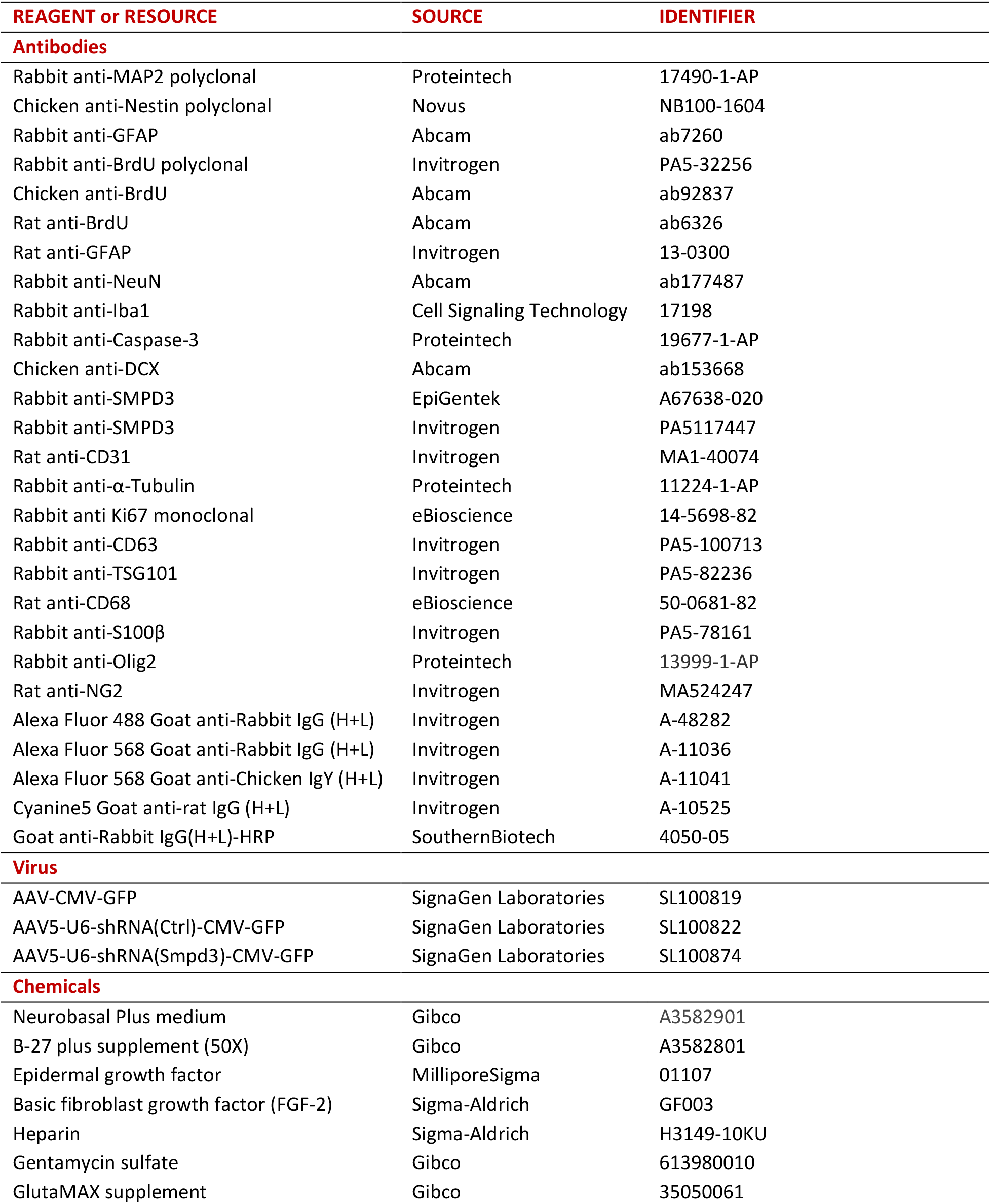
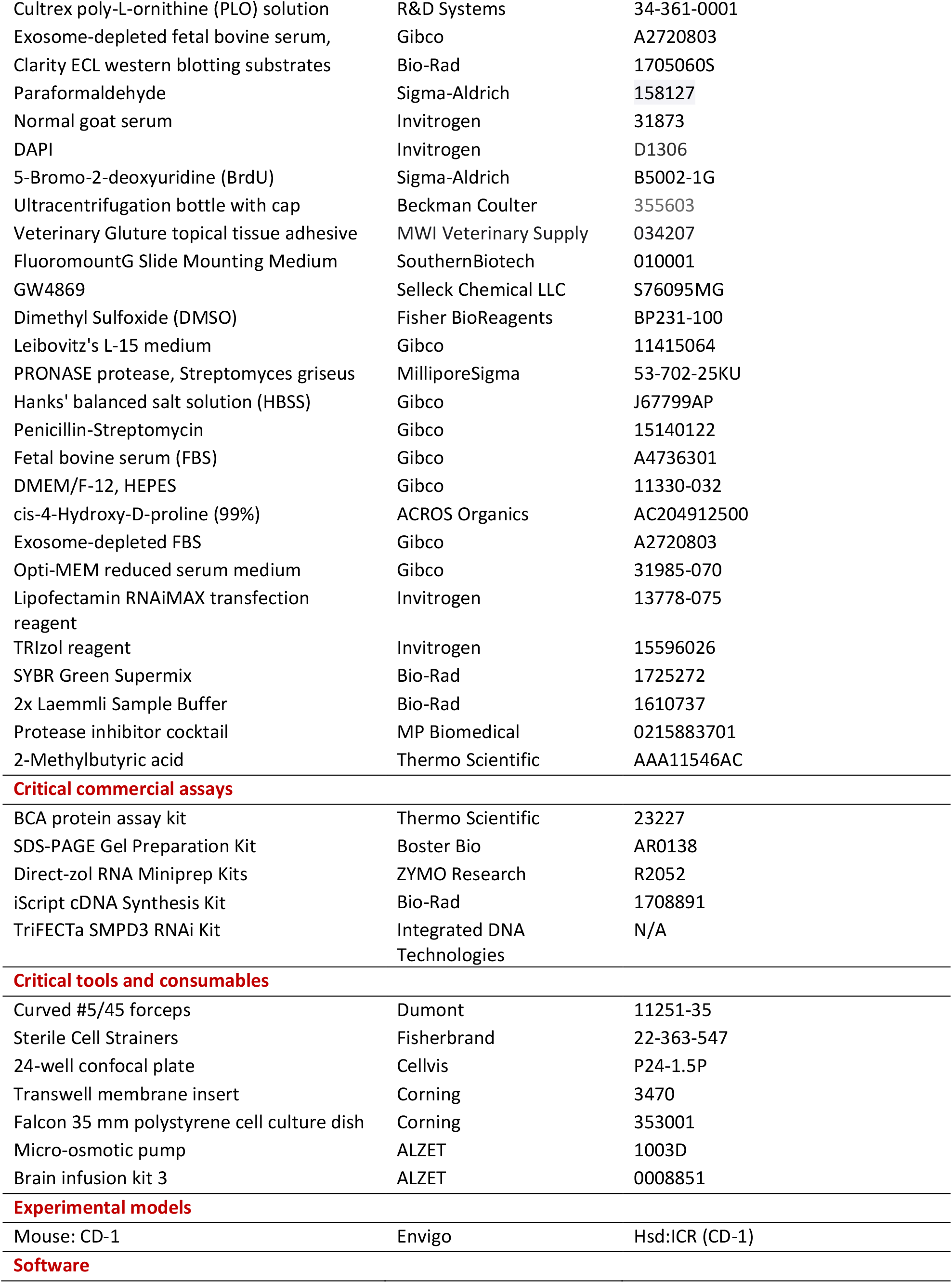
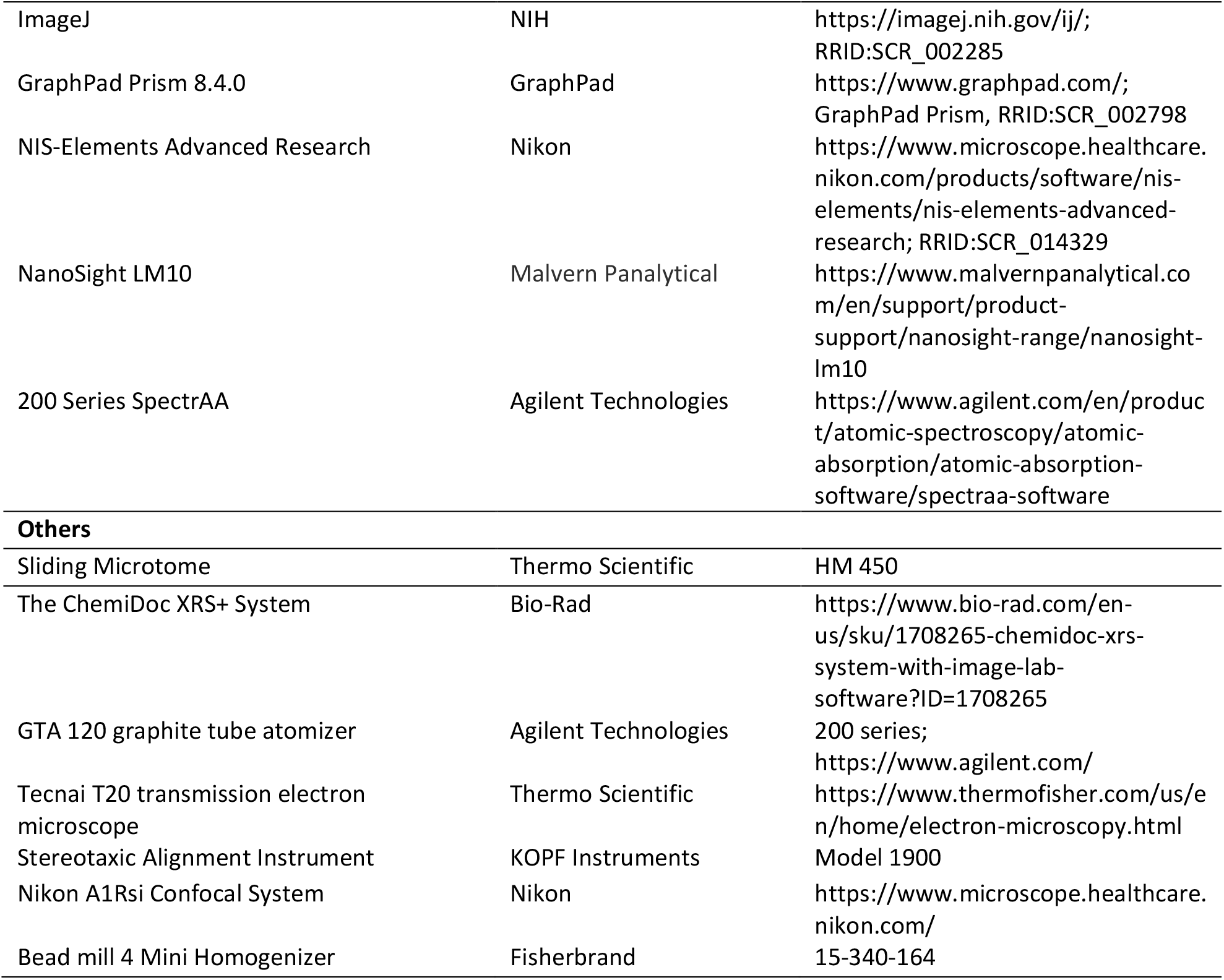
Summary of Resources for Key Chemicals, Reagents and Assays Used in this Report.

### 2.2. Animals

Male CD-1 mice (3 months old) were purchased from Envigo (Indianapolis, IN); the rationale for choosing this sex is that males are more prone to developing olfactory disorders than females (Croy et al. 2014; Noel et al. 2017; Sorokowski et al. 2019; Temmel et al. 2002; Yang and Pinto 2016). Upon arrival, mice were housed in a temperature-controlled, 12/12-h light/dark cycle room and allowed to acclimate for one week prior to experimentation. Animals were provided with deionized water and pellet rat chow (Teklad Dlobal 18% Protein Rodent Diet, 2018s; Envigo) *ad libitum*. This study was conducted in compliance with standard animal use practices and approved by the Animal Care and Use Committee of Purdue University (PUCAC No. 1112000526).

### 2.3. Co-culture of SVZ neurospheres with CP tissues

To investigate the direct impact of CP on SVZ neurogenesis, we established an *in vitro* co-culture system using a two-chamber Transwell device. Freshly isolated CP tissues from brain ventricles were incubated in the upper Transwell insert and the neurospheres derived from adult mouse SVZ tissues were cultured on the glass surface of the lower chamber (Fig. 1A). To produce SVZ-derived neurospheres, the SVZ tissue, which was a thin layer lining the lateral ventricle, was carefully dissected using a pair of Dumont curved #5/45 forceps. Following filtration through a 40-µm cell strainer, the dissociated preparation was cultured in a neurosphere background medium (NBM: Neurobasal plus medium containing 2% B-27, 1% GlutaMAX, and 50 µg/mL gentamicin) supplemented with EGF (20 ng/mL), FGF (20 ng/mL), and heparin (2 µg/mL) for neurosphere induction. Primary neurospheres, which were formed 6 days after the initial seeding, were collected for Trypsin dissociation and the cell suspension was then reseeded into a 96-well U-shaped-bottom microplate (0.1 mL/well, 1.0 × 10^4^ cells/mL in NBM enriched with growth factors and heparin) for secondary neurosphere formation. After 24 hrs, the secondary neurospheres were formed with nearly identical diameters (Liu et al. 2022b). These secondary neurospheres were collected and resuspended in NBM for subsequent plating onto the PLO-coated glass bottom in the lower chamber.

**Figure 1.**
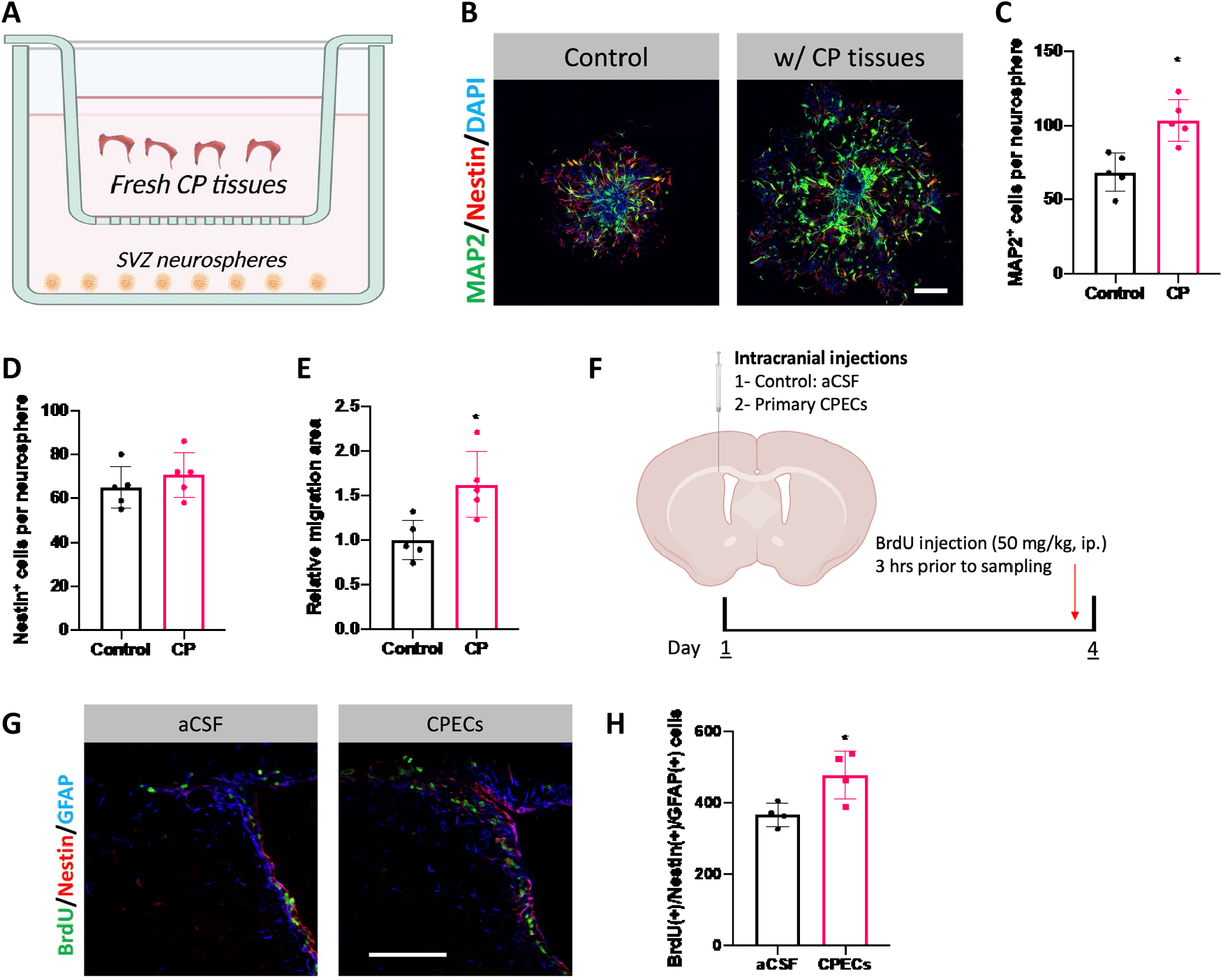
Choroid plexus modulates SVZ adult neurogenesis *in vitro* and *in vivo*. (A) Graphical illustration of CP-SVZ neurosphere co-culture system. (B) Representative immunocytochemistry images demonstrating neurosphere growth affected by CP. Scale bar = 200 µm. (C-E). Quantification of MAP2^+^ neuronal cells and Nestin^+^ progenitor cells per neurosphere, and the relative migration area of neurospheres. n = 5. (F). Schematic illustration of intracranial injection of primary cultured CPECs to the SVZ niche. (G and H). Representative immunohistochemistry images and quantification of proliferating SVZ NSCs following intracranial injection of CPEC. n = 4; scale bar = 100 µm.

CP tissues were dissected from brain ventricles of two PBS-perfused mouse brains. The collected CP tissues were resuspended in 0.6 mL NBM, and then transferred to the Transwell inserts which were placed above the seeded neurospheres. The control group had 0.6 mL NBM only in the insert. Following 24 hrs of co-culture, CP tissues along with the insert were removed. Neurosphere culture continued for additional 6 days. All cell cultures, including this CP-SVZ co-culture, in this study were performed in 95% air/5% CO_2_ at 37 °C. Upon completion of the culture, neurospheres were fixed with 4% PFA in PBS for 10 min under room temperature and stored in PBS until further immunostaining for confocal microscopy analysis.

### 2.4. Immunocytochemistry

Immunocytochemistry was performed as previously described (Liu et al. 2022b) to characterize the neurosphere growth and the impact of co-culture with CP tissues. Fixed neurosphere samples were blocked/permeabilized in PBST containing 5% normal goat serum and 0.3% TritonX-100. Blocked neurospheres were further incubated with primary antibodies against MAP2 (marker for mature neuronal cells) and Nestin (marker for NSPCs) at 4°C overnight. Following 3 washes with PBST, neurospheres were incubated with fluorophore-conjugated secondary antibodies (1:500) at room temperature for 1 h while being protected from light, followed by counterstaining with DAPI. After 3 PBST washes, samples were immediately analyzed utilizing a Nikon A1Rsi confocal system.

### 2.5. Primary culture of choroid plexus epithelial cells (CPECs)

Primary culture of CPECs was performed as previously described (Monnot and Zheng 2012) with modifications. Briefly, freshly isolated CP tissues from 2 mice were washed in Leibovitz’s L-15 medium and then pelleted by centrifugation at 300 g for 3 min. After aspirating L-15 medium, CP tissues were dissociated with pre-warmed pronase (2.5 mg/mL) in HBSS at 37 °C for 7 min. To terminate the digestion, 1 mL CPEC medium, which was prepared by supplementing DMEM/F12 medium with 10% FBS, penicillin (100 U/mL), streptomycin (100 μg/mL), gentamycin (40 μg/mL) and EGF (100 ng/mL), was added, followed by centrifugation (500 g for 5 min) to isolate samples. After supernatant aspiration, CP samples were resuspended in 2-mL CPEC culture medium and further dissociated by pipetting up and down 10 times through a regular P1000 tip. Dissociated cells were then plated on a PLO-precoated 35-mm Petri dish and cultured for 3 days, followed by treatment with cis-HP (25 μg/mL medium) for 3 days to eliminate fibroblast contamination. After a 1-day continued culture with CPEC medium, the primary culture of CPECs was established and ready for the studies described below.

### 2.6. Intracranial injection of primary CPECs

To test whether the CP modulates SVZ NSC activities, primary-cultured CPECs were stereotaxically injected to the SVZ niche, which actively supplies newborn neurons to the OB (Doetsch and Alvarez-Buylla 1996). Since the growth factors such as EGF are known to influence SVZ NSCs (Doetsch et al. 2002), the CPECs were cultured in the medium without EGF and FBS for additional 2 days with daily medium replacement. For brain injections, a suspension of primary CPECs was prepared by incubating primary culture with Trypsin at 37 °C for 3 min. Following 2 wash-centrifugation cycles, CPECs were resuspended in artificial CSF (aCSF) at a density of 1.5 × 10^3^ cells/µL. Upon intracranial injection, mice under isoflurane anesthesia received 1 µL CPEC suspension at the following coordinate: 0.0 mm, 1.5 mm, 2.0 mm (anterior, lateral, ventral distance relative to bregma); the control mice received 1 µL aCSF at the same coordinate. To ensure accurate graft, preparations were slowly injected within 10 min, followed by a 5-min stay before withdrawing the needle from the brain. The incised skin was closed by Gluture topical tissue adhesive alongside the cut, followed by triple antibiotic ointment application. Post-operative animals recovered on a heat pad and were then housed individually for 3 days.

To label proliferating NSCs, BrdU (50 mg/kg) was intraperitoneally (IP) injected 3 hrs prior to necroscopy. Under deep anesthesia by IP injection of ketamine (75 mg/kg) and xylazine (10 mg/kg), mice were sequentially perfused with ice-cold PBS and 4% PFA in PBS. Brain samples were extracted, and immersion fixed overnight with by 4% PFA. Following dehydration in 30% sucrose for 72 hrs, brain samples were mounted on a microtome and frozen by dry ice for slicing. Coronal slices containing the SVZ, which approximately span from +1.5 mm (anterior) to -1.0 mm (posterior) relative to the bregma, were collected from the injected hemisphere (serially into 3 wells for preservation). Slices were preserved in cryopreservation medium (30% sucrose, 1% polyvinylpyrrolidone, 30% ethylene glycol in 0.1 M phosphate buffer) at -20 °C until immunohistochemistry analysis.

### 2.7. Immunohistochemistry

Immunohistochemistry was performed to characterize the SVZ adult neurogenesis as previously described (Liu et al. 2022b). First, brain slices preserved in cryopreservation medium were washed with PBS 3X to remove cryopreservation medium. In subsequent steps, if BrdU analysis was needed, slices were incubated in 2 N HCl for 30 min under 37 °C, followed by neutralization with 0.1 M boric solution (pH 8.5) for 30 min; slices continued to be blocked/permeabilized with PBST containing 5% normal goat serum and 0.3% TritonX-100. Otherwise, PBS-washed slices were directly blocked/permeabilized. Blocked brain slices were then incubated with primary antibodies overnight at 4 °C. Following 3 PBST washes, brain slices were further incubated with fluorophore-conjugated secondary antibodies (1:500) for 1 h at room temperature and then with DAPI for counterstaining (prevented from light). Following 3 PBST washes, brain slices were mounted onto microscope slides with mounting medium. Images were captured by Nikon A1Rsi Confocal system; Z-stack scanning and large-image stitching were coupled for 3D reconstruction and cell counting.

### 2.8. Differential centrifugation of conditioned medium for sEV collection

CP tissues were cultured for 24 hrs in sEV-free CPEC medium, which was prepared by mixing ultracentrifuged DMEM/F12 medium containing penicillin (100 U/mL), streptomycin (100 μg/mL), and gentamycin (40 μg/mL) with ultracentrifuged exosome-depleted FBS (10%). Conditioned medium from primary CPECs were collected at the end of the culture. To begin the differential centrifugation, conditioned medium was centrifuged at 300 g for 10 min to pellet live tissues/cells, followed by a 20-min centrifugation at 2,000 g to remove dead cells/debris. Subsequently, the medium was further centrifuged at 4 °C at 12,000 g for 30 min, leaving large vesicles pelleted for collection. The supernatant medium was then ultracentrifuged at 4 °C at 120,000 g for 70 min, which separated the crude small extracellular vesicles (sEVs) (pellet) from soluble proteins (supernatant). After collecting the soluble proteins, the crude sEV samples were purified by ultracentrifugation for a second time (illustrated in Fig. 2A). Finally, sEVs were resuspended and stored in NBM (for testing neurosphere growth) or PBS (for Western blot, nanoparticle tracking analysis and electron microscopy) at −80 °C.

**Figure 2.**
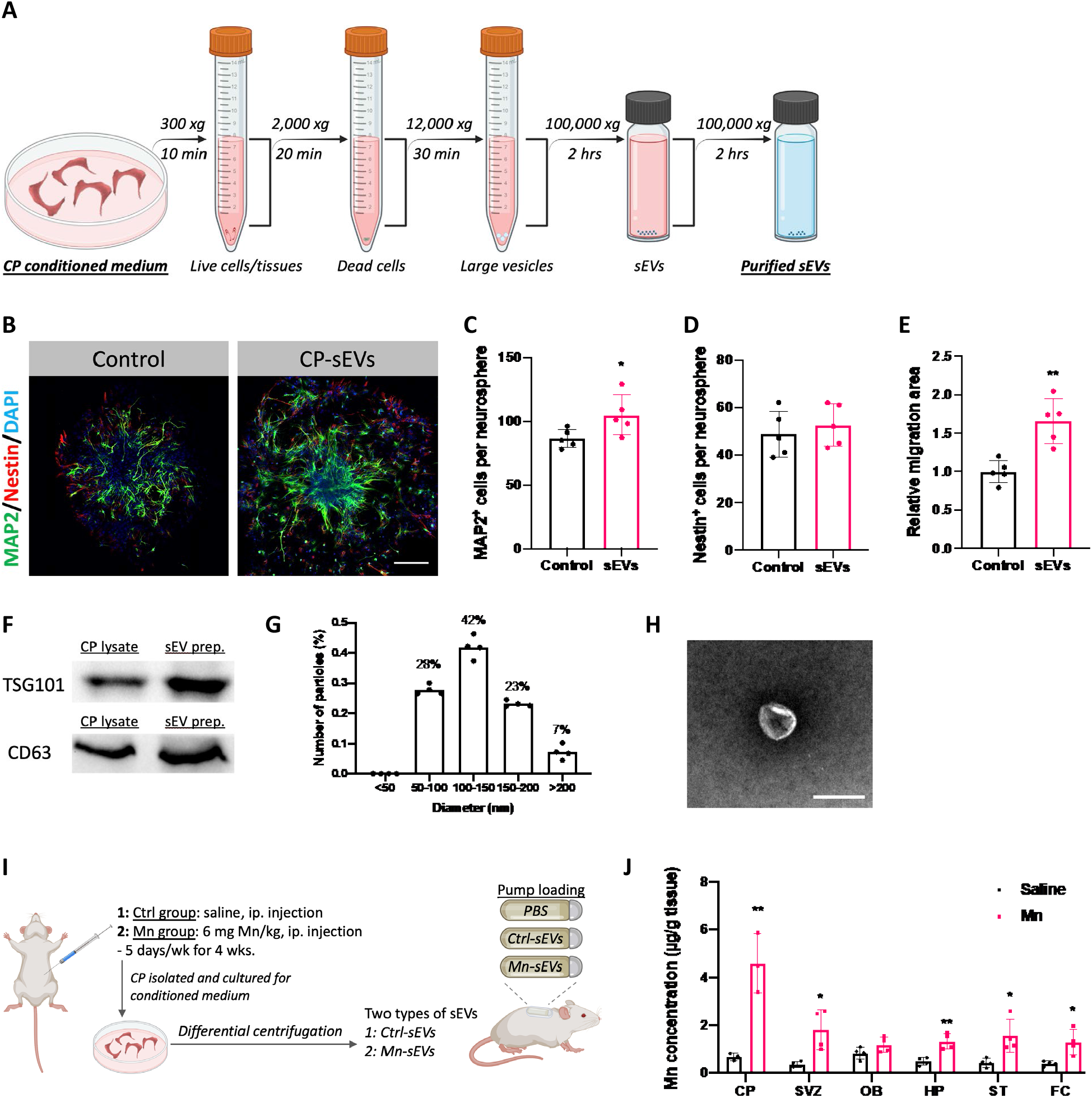
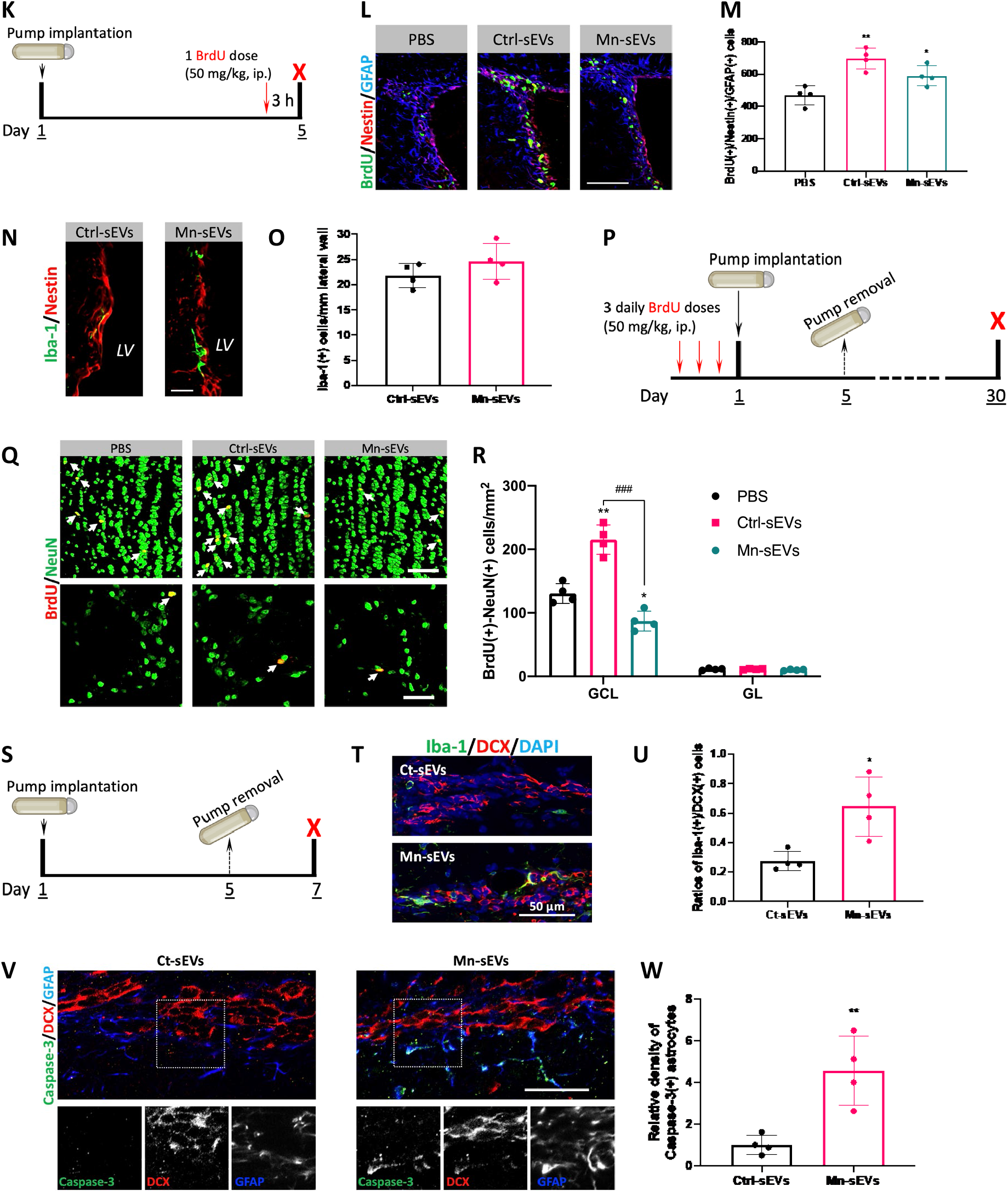
Small extracellular vesicles (sEVs) mediate CP’s regulatory effects on SVZ adult neurogenesis. (A) Diagram showing differential centrifugation of CP conditioned medium for sEV collection. (B) Representative immunocytochemical images showing neurosphere growth as affected CP-sEVs. Scale bar = 100 µm. (C-E). Quantification of MAP2^+^ neuronal cells and Nestin^+^ progenitor cells per neurosphere, and the relative migration area of neurospheres. n = 5. (F). Western blot characterization of CP-secreted sEVs by expression of CD63 and TSG101. (G). Size distribution of CP-secreted sEVs by NTA. n = 4. (H) Electron microscopy image of CP-secreted sEVs. Scale bar = 100 nm. (I) Diagram depicting the experimental design for functional characterization of CP-secreted sEVs in SVZ adult neurogenesis. Control and Mn-treated mice were used to collect sEVs from CP: Ctrl-sEVs and Mn-sEVs. Preparations of CP sEVs were loaded into osmotic pump for ICV infusion, with specific experimental timelines used in (K), (P), and (S) to characterize the SVZ adult neurogenesis. (J) Atomic absorption spectrometry (AAS) analyses to quantify Mn concentrations in multiple brain regions following the 4-week Mn exposure at 6 mg Mn/kg. Tested brain regions included CP, SVZ, OB, hippocampus (HP), striatum (ST) and frontal cortex (FC). n = 3-4. (K) Experimental timeline to study the proliferation of SVZ NSCs following ICV infusion of Ctrl-sEVs and Mn-sEVs (PBS as controls). (L and M). Representative immunohistochemistry images and quantification of proliferating SVZ NSCs as affected by ICV infusions of Ctrl-sEVs and Mn-sEVs. n = 4; scale bar = 100 µm. (N and O). Representative immunohistochemistry images and quantification of Iba-1(+) microglial cells in the SVZ as affected by ICV infusions of Ctrl-sEVs and Mn-sEVs. n = 4; scale bar = 20 µm. (P). Experimental timeline to study the OB newborn neurons as affected by ICV infusions of Ctrl-sEVs and Mn-sEVs (PBS as controls). (Q and R). Representative immunohistochemistry images and quantification of OB newborn neurons as affected by ICV infusions of Ctrl-sEVs and Mn-sEVs. The upper and lower panel in (Q) represents newborn neurons in the OB’s granule cell layer (GCL) and glomerular layer (GL), respectively. n = 4; scale bar = 50 µm. (S). Experimental timeline to assess the RMS as affected by ICV infusions of Ctrl-sEVs and Mn-sEVs (PBS as controls). (T and U). Representative immunohistochemistry images and quantification of microglial cells in the DCX(+) cell-abundant RMS as affected by ICV infusions of Ctrl-sEVs and Mn-sEVs. The microglial alterations in the RMS was presented as the relative density to DCX(+) neuroblasts. n = 4; scale bar = 50 µm. (V and W). Representative immunohistochemistry images and characterization of apoptotic cells in the RMS as affected by ICV infusions of Ctrl-sEVs and Mn-sEVs. Lower panel in (V) magnifies apoptotic signals observed in RMS of the Mn-sEV group. n = 4; scale bar = 40 µm.

### 2.9. Western blot

Western blot was performed as previously described (Liu et al. 2022a). Briefly, proteins were extracted from samples with RIPA buffer supplemented with protease inhibitor. The protein concentrations in samples were determined via a BCA protein assay kit. After mixing with 2× Laemmli sample buffer, samples were heated to 95 °C for 5 min. Upon electrophoresis, 2 µg protein from sEV samples and CP lysates were loaded to verify expression of sEV-related marker (CD63 and TSG101), while the remaining samples analyzed in this study had 10 µg protein loaded into each well. Following electrophoresis under 80 V for 75 min, proteins on the gel were then transferred to PVDF membranes (under 100 V for 1 hr in an ice tank). The PVDF membranes were then blocked with 5% bovine serum albumin in Tris-buffered saline with 0.1% Tween 20 (TBST) and incubated overnight at 4°C with the primary antibodies (1:3000 for α-tubulin and 1:500 for all the other primary antibodies). Following sufficient washes with TBST, PVDF membranes were further incubated with goat anti-rabbit HRP-conjugated secondary antibodies (1:3000) at room temperature for 1 h. The colorimetric signals on the membranes were developed using ECL Western Blotting substrate and captured by ChemiDoc XRS system. The band intensities were quantified using ImageJ and were reported as relative expression levels normalized to α-tubulin.

### 2.10. Nanoparticle tracking analysis (NTA)

NTA was performed as previously described (Hettinger et al. 2021). Briefly, sEV samples diluted in ultracentrifuged sterile-filtered PBS were slowly injected to the imaging chamber of the NanoSight LM10 system. This system was equipped with a 405 nm laser and a sCMOS camera. By setting up camera level at 10 and detection threshold at 4, two 30-sec videos were captured at the “optimal imaging location”, an area just past the flare spot where the laser beam emerges into the chamber. The particle size distribution and concentration of sEV samples were analyzed using the NanoSight NTA software (v2.3).

### 2.11. Electron microscopy

Electron microscopic images of CP-secreted sEVs were acquired under technical guidance by Life Science Microscopy Facility (LSMF) at Purdue University. Briefly, sEV samples were pipetted onto a carbon-coated copper EM grid and incubated for 2 min. After blotting away excess liquid, grids were washed with diH_2_O to remove salts. The grids were then negatively stained with 2% phosphotungstic acid for 1 min. Images were acquired through a Gatan US1000 2K CCD camera on a FEI Tecnai T20 transmission electron microscope equipped with a LaB6 source operated at 200kV.

### 2.12. Manganese (Mn) exposure and atomic absorption spectrometry (AAS)

Mice were exposed to Mn as previously described (Fu et al. 2015, 2016). Briefly, MnCl_2_·4H_2_O (dissolved in sterile saline) was IP injected to mice at the dose of 6 mg Mn/kg, once daily, 5 days/week for 4 consecutive weeks; controls received an equivalent volume of sterile saline. Upon completion of Mn treatment, mice were anesthetized with ketamine/xylazine (75:10 mg/kg, 1 mg/kg, IP) and transcardially perfused with 30 mL ice-cold PBS to remove blood. After extracting the brain, brain tissues (CP, SVZ, OB, hippocampus, striatum, and frontal cortex) were collected for Mn concentration measurements with AAS. Following digestion with concentrated ultrapure HNO_3_ in a MARSXpress microwave-accelerated reaction system, Mn concentrations in brain samples were quantified by 200 Series SpectrAA with a GTA 120 graphite tube atomizer. The detection limits for Mn by this system was 0.09 µg/L. Throughout the measurements, AAS readings were kept in the linear range of the standard curve (0-20 µg/L) by proper dilutions with diH_2_O. Mn concentrations across tested brain regions were reported as “µg Mn/g tissue”.

### 2.13. Intracerebroventricular (ICV) infusions of CP-secreted sEVs using osmotic pumps

With the Mn exposure paradigm, another subset of mice were used to collect sEVs secreted by the CP in control (saline-injected) and Mn-exposed mice. Upon completion of the exposure, CP tissues were isolated from brain ventricles and cultured in sEV-free CPEC medium for 24 hrs. Through differential centrifugation procedures, purified sEVs were acquired from the CP conditioned medium: Ctrl-sEVs (from the saline controls) and Mn-sEVs (from the Mn-exposed mice). Resuspended in PBS, sEV samples were immediately loaded into osmotic pumps for the 4-day ICV infusions (PBS only as infusion controls); each pump contained sEVs isolated from CP tissues of 2 mouse brains (illustrated in Fig. 2I). The stability of CP-sEVs was assessed by incubating such samples in a 37 °C incubator for 4 days and monitoring the sEV concentrations by NTA. For different experimental purposes, BrdU was administered accordingly (Fig. 2K and 2P): one IP dose 3 hr prior to sacrifice to label proliferating cells in the SVZ or 3 daily preceding doses prior to surgery to trace newborn neurons in the OB. Procedures for ICV infusion were detailed previously (Liu et al. 2022b). Briefly, loaded pumps were coupled with Alzet brain infusion kit 3 to establish a pump-tubing-cannulation system. Isoflurane-anesthetized mice were fixed onto a stereotaxic device. After exposing and clearing the bregma area on the skull, an extended incision was sagittally made to the neck area. Subsequently, a sterile curved hemostat was inserted subcutaneously (curved side up) and slowly advanced to the back area; the hemostat was gently opened and closed to create a space for pump implantation. A burr hold was then drilled at the following coordinate: 0.22 mm, 1.0 mm (posterior, lateral relative to bregma). The pump was then gently inserted into this newly created subcutaneous space, with the cannulation piece (connected with the tubing) available for “pick-up” for the cannula holder. Clamped by the holder, the cannula unit (with Loctite applied on its base) was slowly lowered down to let the steel cannula (2.5 mm) into the brain through the burr hole by 2.5 mm relative to the skull surface and was thus tightly glued. The incised skin was closed by Gluture topical tissue adhesive alongside the cut, followed by triple antibiotic ointment application. The postoperative animals were recovered on a heat pad and housed separately. Upon completion of the ICV infusion, mice were either sacrificed for sampling (Fig. 2K) or had pump removed (under anesthesia) followed by closure of tubing’s free end and survived until the planned sampling date (Fig. 2P and 2S). Upon completion of ICV infusion experiments, brain sampling (perfusion, fixation, and dehydration) and microtome slice preparation (serially into 6 wells) were performed as described before. Brain slices were stained through immunohistochemistry to characterize the SVZ adult neurogenesis as affected by ICV infusions.

### 2.14. Chemical inhibition of sEV secretion

A noncompetitive inhibitor of sphingomyelinase, GW4869, was IP administered as previously described to inhibit sEV secretion (Essandoh et al. 2015). Briefly, GW4869 was dissolved in DMSO and IP injected for 3 consecutive days at the dose of 2.5 mg/kg; the controls injected with equivalent amount of DMSO. Six hrs after the last GW4869 dose, mice underwent CSF collection and brain tissues dissection. Collected CSF samples, following differential centrifugation, were analyzed with NTA to quantify sEV concentrations. Dissected SVZ tissues were used for qPCR to quantify transthyretin mRNA (Ttr) level with or without GW4869 treatment (Fig. 3A).

**Figure 3.**
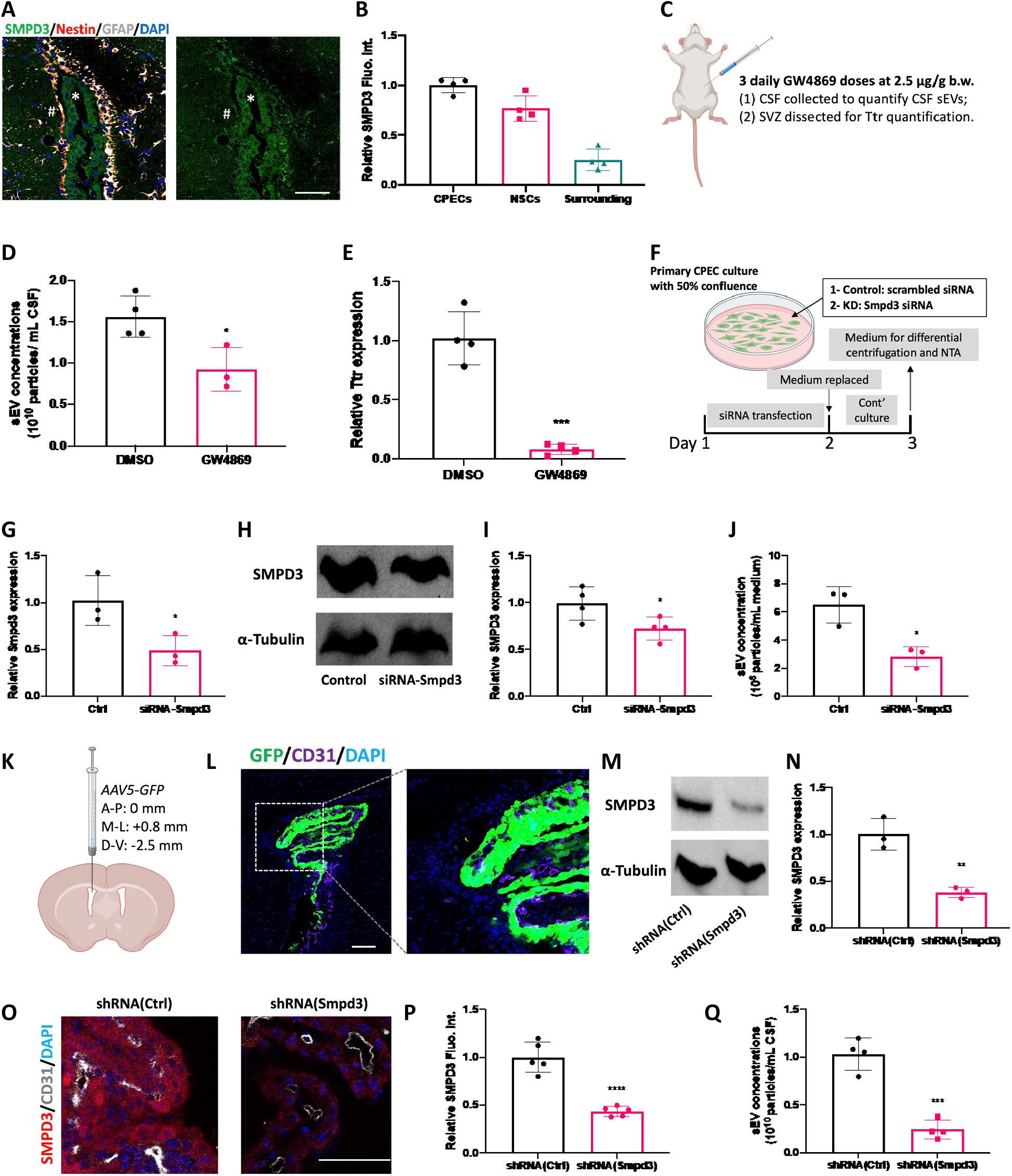
Secretion of sEVs by the CP is determined by SMPD3. (A and B). Representative immunohistochemistry image showing SMPD3 expression in the CP, SVZ, and surrounding and quantitative analysis of SMPD3 immunofluorescence intensity in these regions. n = 4; scale bar = 50 µm. (C) Experimental design for chemical inhibition of SMDP3 by GW4869 administration. (D) Quantification of sEVs in the CSF by NTA as affected by GW4869 administration. n = 4. (E). Relative Ttr level in the SVZ by qPCR as affected by GW4869 administration. n = 4. (F) Experimental timeline to study the role of SMDP3 in sEV secretion by siRNA transfection using primary-cultured CPECs. (G) Relative Smpd3 mRNA level in CPECs following siRNA transfection by qPCR. n = 4. (H and I). Representative immunoblotting images and quantitative analysis of SMPD3 protein expression in CPECs following siRNA transfection. n = 4. (J) NTA of sEV concentration in CPEC conditioned medium following siRNA transfection. n = 4. (K). Diagram showing ICV injections of AAV5 to the lateral ventricle. (L). Immunohistochemistry image showing AAV5 tropism toward CPECs of the CP. Scale bar = 100 µm. (M and N). Representative immunoblotting images and quantitative analysis of SMPD3 protein expression in the CP tissues following CP-targeted SMPD3 knockdown by AAV5. n = 3. (O and P). Representative immunohistochemistry images and quantitative analysis of SMPD3 in the CPECs following CP-targeted SMDP3 knockdown. n = 5; scale bar = 50 µm. (Q). NTA of sEV concentration in the CSF following CP-targeted SMDP3 knockdown. n = 4.

### 2.15. CSF collection

The CSF samples were collected by cisterna magna puncture with a fire-pulled glass pipette. Briefly, under isoflurane anesthesia, the head of mice was fixed onto a stereotaxic device. By sequentially trimming the fur on the neck, sagittally incising the skin, removing the connective tissues, and severing 3 covering muscular layers, dura mater with a pale/tough texture was spotted. The fire-pulled pipette was carefully advanced, while avoiding evident blood vessels, for cisterna magna puncture. CSF samples in the pipette were carefully released into Eppendorf tubes. By a brief centrifugation at 2,000 g for 5 min, CSF samples were inspected for any blood contamination (evident red spot on the bottom); contaminated CSF samples were discarded.

### 2.16. Quantitative polymerase chain reaction (qPCR)

qPCR was performed as previously described (Liu et al. 2022b). Briefly, total RNA was extracted from samples using TRIzol reagent and Direct-zol RNA Microprep kit. Extracted RNA (0.5 µg/per sample) was reverse transcribed into cDNA using BioRad iScript cDNA synthesis kit. Mixing cDNAs with specific primers and SYBR Green Supermix, the reactions occurred under the following programmed heating cycles for CT values: initial 3 min denaturation at 95 °C, followed by 40 cycles of 30 s denaturation at 95 °C, 10 s gradient 55.0–65.0 °C, and a 30 s extension at 72 °C. Dissociation curves were examined to verify that the majority of detected fluorescence was derived from the labeling of specific PCR products. Each qPCR reaction was run in duplicate. Relative mRNA expressions were calculated using 2^−∆∆Ct^ method with beta-actin (Actb) as the reference gene. The forward and reverse primers for genes of interest in this study were designed by Primer Express 3.0 software and listed in Table S2.

**Table S2.**
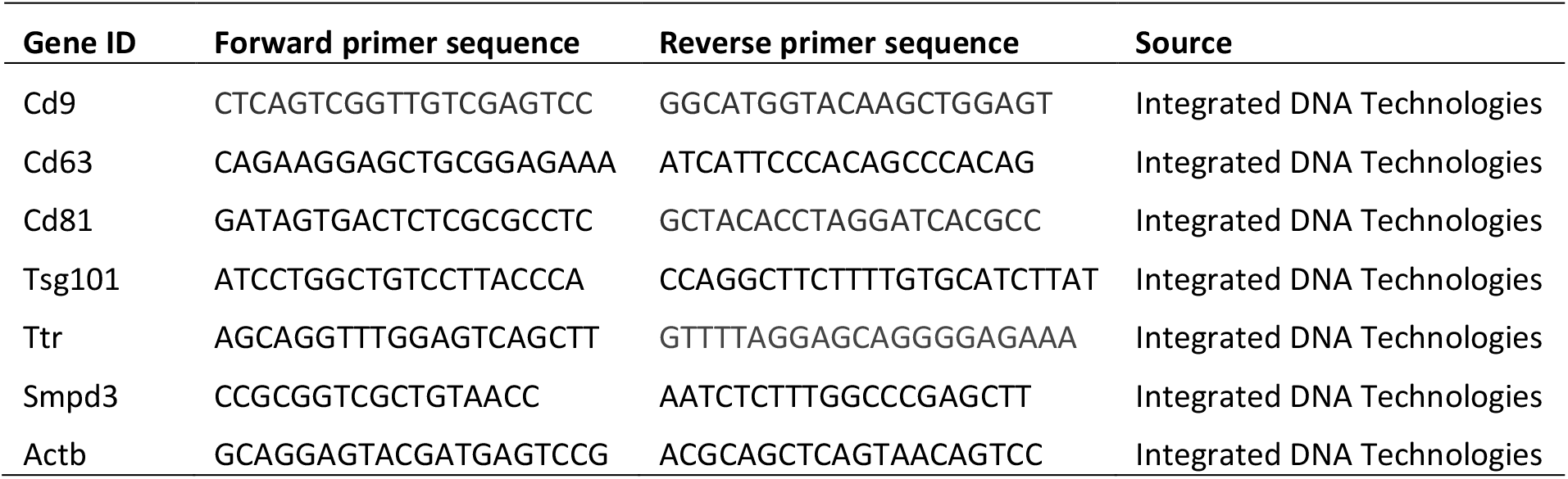
qPCR primers used in this study.

### 2.17. siRNA knockdown

Primary CPECs were grown to a confluence of approximately 50% in CPEC culture medium. To transfect siRNA, culture medium was removed, and cells were washed with Opti-MEM medium for a serum-free environment. For SMPD3 knockdown, Opti-MEM medium containing Smpd3-targeting siRNA (10 nM) and 0.5% Lipofectamin RNAiMAX was prepared, mixed, and added to the culture; controls were cultured with medium containing nontargeting siRNA (otherwise the same). Cells were incubated with siRNA for 48 hrs. In first experiment to assess the siRNA transfection efficacy, cells were then collected to quantify Smpd3 mRNA level by qPCR and SMPD3 protein expression by Western blot. In the second experiment to determine the role of SMPD3 in CPEC sEV secretion, following the 48-hr transfection, culture medium was replaced with sEV-free CPEC medium for an extended culture of 1 day. The culture medium was then collected for differential centrifugation for sEV concentration measurements by NTA.

### 2.18. Intracerebroventricular (ICV) injections of AAV5

The AAV5 used in this study was manufactured by SignaGen Laborotories. In testing viral tropism toward CP, AAV5-CMV-GFP was used. To selectively knockdown SMPD3 expression in the CP, AAV5-U6-shRNA(mSmpd3)-CMV-GFP was used, with the scrambled sequence-carrying AAV5-U6-shRNA(SCRM)-CMV-GFP as controls; these 2 shRNA-carrying AAV5 vectors were hereafter abbreviated as AAV5-shRNA(Smpd3) and AAV5-shRNA(Ctrl), respectively. For different purposes, BrdU was administered through IP injections accordingly (Fig. 4A, 4D, and 4G): one dose 3 hrs preceding sacrifice to label proliferating cells or 3 daily doses one month prior to sampling to trace newborn neurons in the OB. For AAV5 injections, isoflurane-anesthetized mice were fixed onto a stereotaxic frame. With a sagittal incision, the bregma was exposed and then made clear by applying 30% hydrogen peroxide. A cranial burr hole was then drilled on the skull on the left hemisphere with the following coordinates: 0.8 mm, 0.0 mm (lateral and anterior relative to bregma). Subsequently, a 30-gauge needle of a 10-µL Hamilton syringe prefilled with 3 µL AAV5 solution (4 × 10^12^ VG/mL) was lowered down through the burr hole by 2.5 mm to reach the lateral ventricle. The AAV5 solution was then slowly injected to the CSF at a rate of 0.5 µL/min over 6 min; the needle remained unmoved in the ventricle for an additional 5 min before withdrawal. The incision was then closed by topical tissue adhesive followed by triple antibiotic ointment application. Post-operative animals were recovered on a heating pad and housed individually thereafter. Upon completion of the designed experimental timelines, brain sampling (perfusion, fixation, and dehydration) and slice preparation (serially into 6 wells) were conducted as described previously. Immunohistochemistry was performed with brain slices to study the effects of AAV5 ICV injection on the SVZ adult neurogenesis.

**Figure 4.**
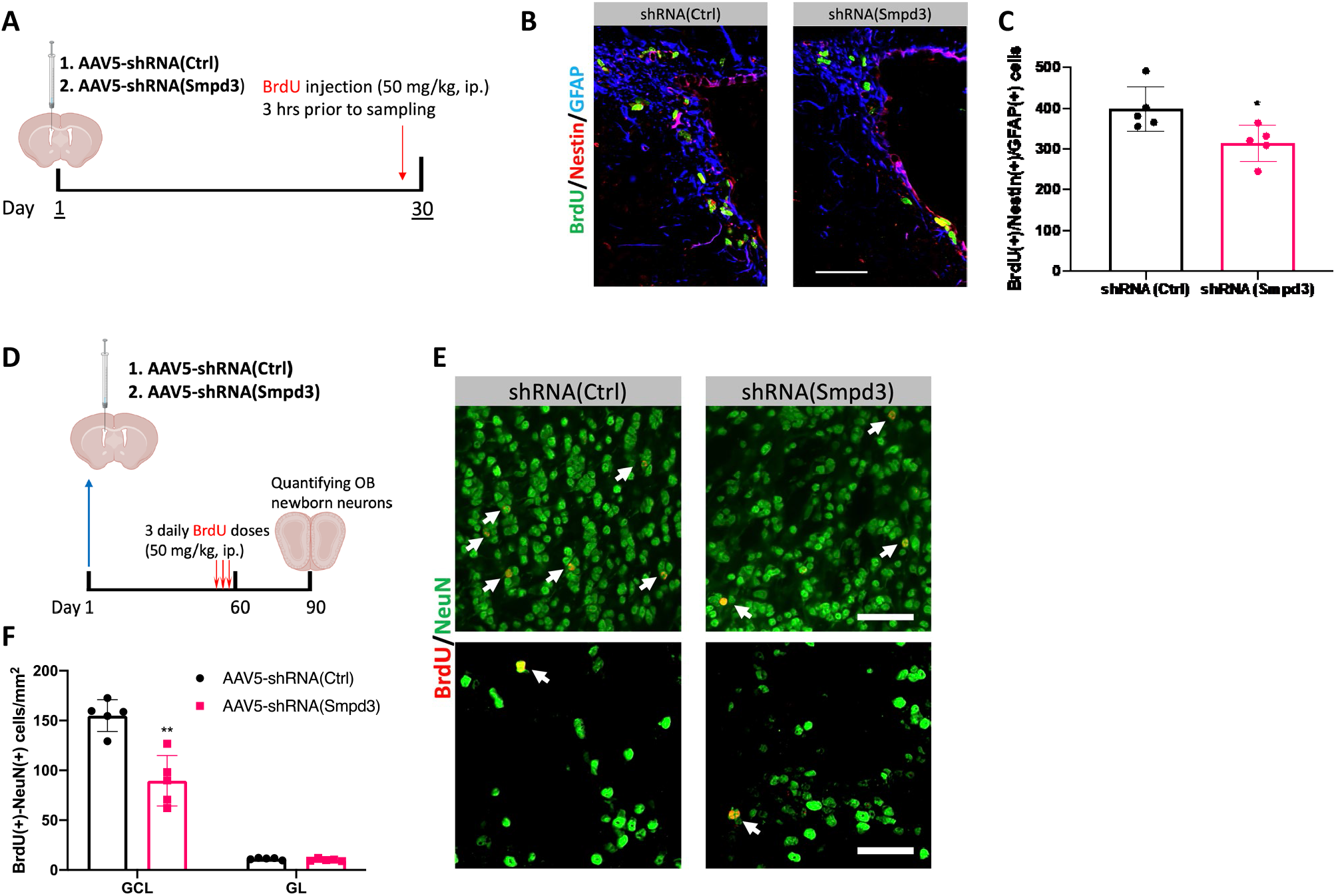
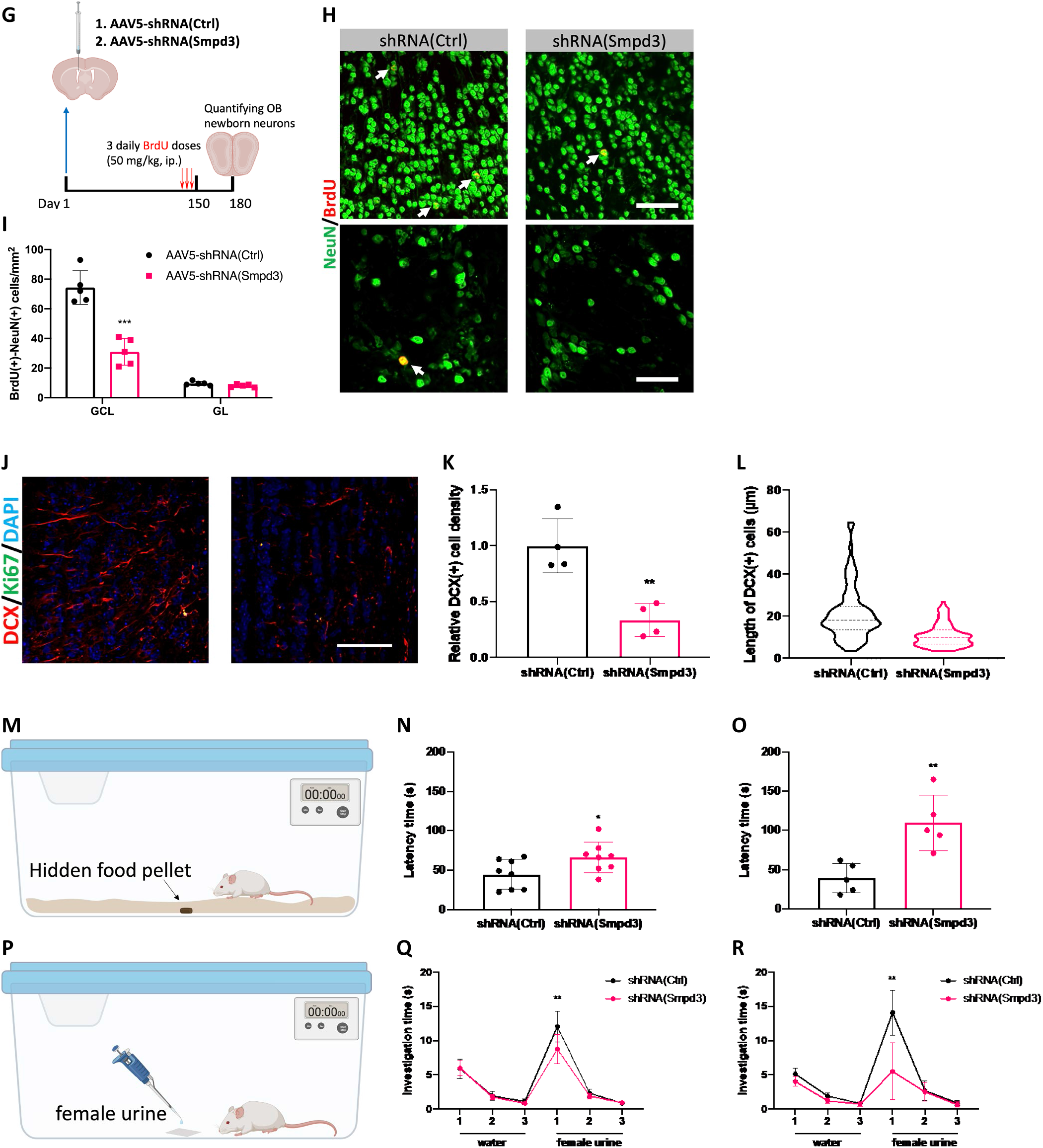
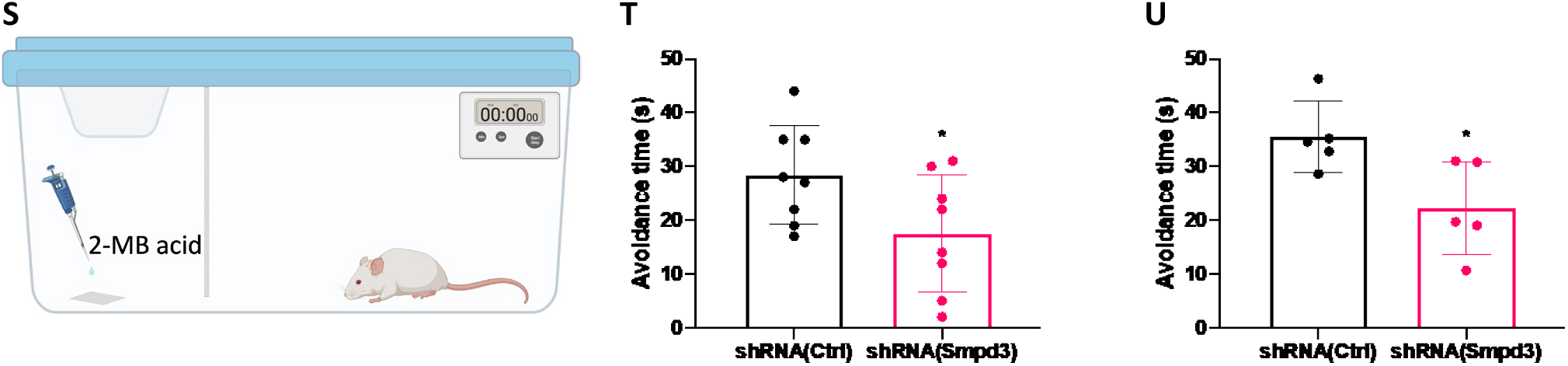
CP-targeted sEV secretion suppression caused diminished SVZ adult neurogenesis and compromised olfactory performance. (A). Experimental timeline to study proliferation of SVZ NSCs as affected CP-targeted sEV secretion suppression. (B and C). Representative immunohistochemistry images and quantification of proliferating SVZ NSCs. N = 5; scale bar = 50 µm. (D). Experimental timeline to study newborn neuron generations as affected by short-term CP sEV secretion suppression. (E and F). Representative immunohistochemistry images and quantification of OB newborn neurons as affected by short-term CP sEV secretion suppression. Upper and lower panels in (E) represent newborn neurons in the GCL and GL, respectively. n = 5; scale bar = 50 µm. (G). Experimental timeline to study newborn neuron generations as affected by long-term CP sEV secretion suppression. (H and I). Representative immunohistochemistry images and quantification of OB newborn neurons as affected by long-term CP sEV secretion suppression. Upper and lower panels in (H) represent newborn neurons in the GCL and GL, respectively. n = 5; scale bar = 50 µm. (J, K, and L). Representative immunohistochemistry images, density quantification, and length measurements of DCX(+) cells in the OB as affected by long-term CP sEV secretion suppression. n = 4; scale bar = 50 µm. (M). Diagram showing the pellet-finding assay. (N and O). Quantitative analysis of the latency time to find the hidden pellet as affected by short- and long-term CP sEV secretion suppression. n = 8 for short-term and n = 5 for long-term assessment. (P). Diagram showing the olfactory preference test. (Q and R). Quantitative analysis of investigation time of female urine as affected by short- and long-term CP sEV secretion suppression. n = 8 for short-term and n = 5 for long-term assessment. (S). Diagram showing the olfactory avoidance test. (T and U). Quantitative analysis of the avoidance time to 2-MB acid presentation in the larger compartment relative to diH_2_O exploring time as affected by short- and long-term CP sEV secretion suppression. n = 8 for short-term and n = 5 for long-term assessment.

### 2.19. Buried Food Test

The buried food test was performed as described with modifications (Yang and Crawley 2009). Prior to testing, a 24-hr food deprivation was conducted by transferring the mice to new cages with water but without food pellets on the wire top. Testing cages were prepared by filling clean cages with 560 g Teklad laboratory animal corncob bedding, which, following smoothing to flat, creates a 3.5-cm layer. One piece of food pellet weighing 2 g was then hidden underneath the corncob at the center of the testing cage. Before the pellet-finding test, mice were habituated in clean cages (without pellet) for 15 min. Subsequently, mice were transferred from habituation cages to testing cages: a digital stopwatch was used to count the latency time (s) of mice finding the hidden pellet. The “finding” event was defined by observing the pellet unearthed by mice from the 3.5-cm corncob bedding. Of note, the food pellet was hidden underneath at the center but not the corners of cages; this change from others (Yang and Crawley 2009) was based on our observation that CD-1 mice, upon transfer to a new cage, tend to explore the cage corners and edges by digging the bedding materials in the beginning seconds, which could confuse the experimenter in defining the “finding” event.

### 2.20. Olfactory preference test

This olfactory preference test was performed in the form of habituation/dishabituation as previously described (Kobayakawa et al. 2007). Prior to testing, a mouse was placed in the testing cage to acclimate for 15 min. To start a trial, a piece of clean filter paper (1 cm x 1 cm) was adhered to the center of the testing cage followed by 20 µL water pipetted onto this paper. In the following 2 min, the cumulative time spent sniffing the filter paper (investigation time) was recorded with a digital stopwatch. This presentation of water was repeated twice with a 1-min interval (habituation). On the fourth trial (dishabituation), female urine was introduced to the subject by pipetting 20 µL urine sample onto the filter paper; this was also repeated twice with a 1-min interval. Investigation time in the six 2-minute sessions was plotted to assess the subjects’ preference toward female urine as affected by CP-targeted sEV secretion suppression.

### 2.21. Olfactory avoidance test

In the olfactory avoidance test, 2-MB acid (98.0% purity, 9.16 M), a proven olfactory repellent, was used as previously reported (Kobayakawa et al. 2007). In this test, the testing cage was divided into two compartments (1:2) by a partition curtain, which was made by a print paper with one end cut into strips; animals can freely cross compartments through this curtain. Upon testing, a mouse was first habituated to this cage for 15 min. Subsequently, a filter paper pipetted with 20 µL diH_2_O was introduced into the smaller compartment. During the following 2-min session, the time spent (s) in the larger compartment (avoidance time for diH_2_O) was recorded by a digital stopwatch. In the following 2-min session, after removing the diH_2_O filter paper, 2-MB acid solution (1,000x diluted with diH_2_O, 20 µL) was introduced the smaller compartment. Likewise, the time spent in the larger compartment was recorded (avoidance time for 2-MB acid). Ultimately, the avoidance behavior was assessed by the “avoidance time for 2-MB acid” subtracted by the “avoidance time for diH_2_O”.

### 2.22. Statistical analysis

Data in this study are presented as mean ± standard deviation (S.D.). Unpaired two-tailed Student’s t test was used for detecting difference between two groups; for comparisons among multiple groups, we used one-way ANOVA followed by Tukey’s post hoc multiple comparisons test. Graphs and statistical analyses were performed with GraphPad Prism version 8.0. The *p*-values in this report were designated as \**p* < 0.05, \*\**p* < 0.01, \*\*\**p* < 0.001, \*\*\*\**p* < 0.0001 (when compared with controls), ### *p* < 0.001 (for ANOVA post hoc comparisons between non-control groups) and non-significant *p* > 0.05.

## 3. Results

### 3.1. In vitro and in vivo evidence to support the modulation of SVZ adult neurogenesis by CP

To establish the relationship between CP and adult neurogenesis activities in SVZ, a two-chamber co-culture system to mimic the CP and SVZ interaction was utilized, i.e., placing freshly dissected CP tissues in the insert above the SVZ neurospheres cultured on the lower chamber (Fig. 1A). Co-culture of CP with SVZ neurospheres for 24 hrs, as the isolated CP tissues remained largely viable within the first 24 hrs *in vitro*, resulted in a significantly enriched growth of SVZ neurospheres in comparison with controls (Fig. 1B). MAP2(+) cells per neurosphere were increased by 51.1% when co-cultured with CP (68.4 ± 12.9 in controls vs.103.4 ± 14.1 with CP; *p* < 0.05) (Fig. 1C). The presence of CP also promoted the neurosphere migration by 62.4% (*p* < 0.05) (Fig. 1E). However, the Nestin(+) progenitor cells did not differ between the two groups (Fig. 1D). Thus, our initial *in vitro* co-culture data demonstrated that CP promoted the SVZ adult neurogenesis by stimulating the neuronal differentiation and migration without evidently affecting the progenitor cells.

To determine if the CP exerted a similar stimulating effect on SVZ NSCs *in vivo*, primary-cultured CPECs were collected and intracranially injected into the SVZ niche area where newborn neurons are produced for OB (Doetsch and Alvarez-Buylla 1996) (Fig. 1F). Compared with aCSF-injected controls, injection of CPECs significantly elevated the proliferation of GFAP(+)/Nestin(+) NSPCs in the SVZ by 30.2% (Fig. 1G,H), suggesting that some yet-unidentified factors released from CPECs prompted SVZ NSPCs into a pro-neurogenesis status. Hence, we sought to identify the critical secreted extracellular fraction by CP that exerted the NSPC-stimulating properties.

### 3.2. Small extracellular vesicles (sEVs) are the critical CP-secreted mediator to regulate SVZ adult neurogenesis

The sEVs have been recognized as an effective means for intercellular communication (Corrado et al. 2013; Gurung et al. 2021; György et al. 2015). Thus, we set out to assess the ability of the CP to produce and secrete sEVs by comparatively analyzing sEV marker expression across brain regions, including the CP, SVZ, striatum, hippocampus, and cerebellum (Fig. S1). Our data revealed that transcriptions of sEV markers, including Cd9, Cd63, Cd81, and Tsg101, were elevated in the CP in contrast to other evaluated regions (Fig. S1A-S1D). For example, Cd9 mRNA in the CP was 77.0% higher than all the other tested regions (Fig. S1A) and Cd63 was 90.3% higher in the CP than in other tested regions (Fig. S1B). Although the enriched expressions in the CP were somewhat less for Cd81 and Tsg101, the CP still had the highest expressions of both (Fig. S1C and S1D). These qPCR data were further verified at the protein level. Immunohistochemical staining demonstrated that CD63 (Fig. S1E) and TSG101 (Fig. S1E) were enriched in the CP, particularly the CPECs, the cell type in direct contact with CSF for substance secretion. Therefore, our observations corroborate the presence of sEVs in the CP, prompting us to examine if CP-secreted sEVs mediate SVZ adult neurogenesis.

Using differential centrifugation, we acquired the sEVs from the CP secretome (Fig. 2A). By supplementing sEVs to SVZ neurosphere culture medium, our immunofluorescent staining data revealed sEVs significantly enriched neurosphere growth (Fig. 2B) in that neuronal differentiation and migration increased by 21.3% and 65.6%, respectively (Fig. 2C and 2E). To further verify the biological identity of these sEVs, we performed Western blot, nanoparticle tracking analysis (NTA), and electron microscopy, which are necessary for characterization of sEVs as recommended by the International Society of Extracellular Vesicles (Théry et al. 2018). The Western blot analysis demonstrated these sEVs were rich in sEV-specific markers TSG101 and CD63 (Fig. 2F). NTA for size distribution analysis confirmed the diameters of capture particles were primarily smaller than 200 nm (Fig. 2G). Importantly, under electron microscopy, sEVs exhibited the typical “cup-shaped” morphology with diameter less than 200 nm. These experimental approaches confirmed that the sEVs collected from CP in this study met the criteria of exosomes.

To understand the relationship between CP-secreted sEVs and SVZ adult neurogenesis *in vivo*, we characterized the neurogenic process following ICV infusion of CP-derived sEVs in mice. Additionally, we used Mn as a model toxicant to observe if the sEVs collected from the CP of Mn-exposed mice differentially affected adult neurogenesis, compared to the control sEVs from healthy CP. Thus, a group of mice received daily IP injections of Mn at 6 mg/kg for 4 wks; controls received saline. The fractions of CP-derived sEVs were isolated after differential centrifugation (defined as “Mn-sEVs” and “Ctrl-sEVs” for Mn-exposed and control mice, respectively), and loaded into the osmotic pump for ICV infusion (Fig. 2I). Metal analysis by atomic absorption spectrometry (AAS) verified that subchronic Mn exposure increased Mn accumulation in the CP much greater than in other tested brain regions (Fig. 2J). We designed 3 experimental timelines to characterize the impact of CP-secreted sEVs (control vs. Mn-exposed) on NSPC proliferation in SVZ (Fig. 2K), newborn neuron generation in the OB (Fig. 2P), and neuroblast migration in RMS (Fig. 2S).

First, a 4-day infusion of CP-derived sEVs, regardless of the host’s exposure status, significantly increased the proliferating NSPCs in the SVZ (Fig. 2L and 2M). Compared to controls with PBS infusions, Ctrl-sEVs induced a greater proliferation of SVZ NSPCs (by 48.7%) than did Mn-sEVs (by 25.6%) (Fig. 2M). The data suggest that SVZ NSPCs were highly sensitive to CP-secreted sEVs. Although Mn is known to induce microgliosis in the SVZ (Fu et al. 2015), our current data showed that ICV infusion of Mn-sEVs did not significantly affect the microglial density in the SVZ (Fig. 2N and 2O).

Second, to study newborn neuron supply to the OB, Ctrl-sEVs and Mn-sEVs were infused for 4 days and, on day 30, OB tissues were dissected to quantify the newborn neuron density in both granule cell layer (GCL) and glomerular layer (GL) (Fig. 2P). Compared with PBS-infused controls, infusion of Ctrl-sEVs significantly increased the newborn neuron density in the OB’s GCL by 65.4%, while Mn-sEVs evidently decreased this density by 33.1% (p<0.05) (Fig. 2Q and 2R). ICV infusion of CP sEVs, however, did not change the density of newborn neurons in GL (Fig. 2Q and 2R). These findings indicated that normal CP-sEVs played a critical role in maintaining the newborn neuron supply to the OB; Mn accumulation in the CP, however, seemed likely to alter CP-secreted sEVs.

Finally, to understand the discrepancy between increased proliferation in SVZ and diminished newborn neurons in OB following Mn-sEV infusion, we investigated the neuroblast migration in the RMS (Fig.2S). Our data revealed that, in the initial segment of RMS (the portion in proximity to brain ventricle), infusion of Mn-sEVs indeed led to much more microglial cells than did Ctrl-sEVs, as shown by an escalated ratio of Iba-1(+)/DCX(+) cells (Fig. 2T,U). Noticeably also, in Mn-sEV infusion group, Iba-1(+) microglial cells displayed an enlarged body size in both SVZ and the initial RMS segment, indicative of an activated (phagocytotic) phenotype (Fig. 2N and 2T). In addition, we observed a significantly elevated apoptosis along the RMS in Mn-sEV group, in which the apoptotic signal was mainly present in the RMS-supporting astrocytes but not in the DCX(+) neuroblasts (Fig. 2V and 2W). Taken together, these findings demonstrated that CP-secreted sEVs were indeed potent modulators for the SVZ adult neurogenesis.

### 3.3. SMPD3 determines sEV secretion by CPECs

Our next step focused on the molecular pathways in the CP modulating sEV secretion. SMPD3 (Sphingomyelin phosphodiesterase 3) catalyzes the biogenesis and release of sEVs into extracellular spaces (Choezom and Gross 2022; Kosaka et al. 2013; Menck et al. 2017; Poggio et al. 2019). Our immunohistochemical staining data demonstrated the presence of SMPD3 in the CP; moreover, its expression was much higher in CPECs and SVZ NSPCs than in surrounding brain regions (Fig. 3A and 3B). GW4869, a chemical known to noncompetitively inhibit SMPD3 enzymatic activity and disrupt sEV biogenesis/release, has been proven highly effective in alleviating inflammation-induced adverse outcomes by blocking sEV-dependent inflammatory transmission in multiple organs including the brain (Balusu et al. 2016; Essandoh et al. 2015; Lepko et al. 2019). GW4869 was utilized to examine the relationship between SMPD3 activity and sEV release from the CP. Mice received 3 daily IP injections of GW4869 (2.5 µg/g) or equivalent volumes of DMSO (as controls) (Fig. 3C). Following the GW4869 administration, the level of sEVs in the CSF, determined by differential centrifugation and NTA, was significantly lower in GW4869-treated animals than in controls (Fig. 3D). Our qPCR data also showed that the level of Ttr in the SVZ, which was highly enriched in CP (Lein et al. 2007) and stored in sEVs (Lee et al. 2019), was significantly diminished by GW4869 treatment (Fig. 3E).

To further verify the critical role of SMPD3 in regulating sEV secretion by the CP, Smpd3 expression in primary cultured CPECs was knocked down via siRNA transfection (Fig. 3F). Transfection of siRNAs targeting Smpd3 significantly reduced the mRNA expression of Smpd3 in the primary CPECs (Fig. 3G) as well as significantly decreasing SMPD3 protein levels (Fig. 3H and 3I). Importantly, the diminished SMPD3 expression led to suppression of sEV secretion by CPECs (Fig. 3J). These *in vitro* data demonstrated SMPD3 regulated the sEV secretion by CPECs.

We then extended our *in vitro* findings by selectively knocking down SMPD3 in the choroid plexus of an *in vivo* mouse model. Adenovirus-associated virus type 5 (AAV5) can selectively target CP for genetic modifications through ICV injections (Chen et al. 2020; Jang and Lehtinen 2022). By ICV injection of AAV5 and analysis 2 weeks post-treatment (Fig. 3K), our triple staining with GFP (transgene carried by AAV5), CD31 (marker for CP endothelium), and DAPI (nuclear marker) revealed that AAV5 possessed a unique tropism toward the CSF-contacting CPECs, with only minimal expression in CP endothelium (Fig. 3L). More importantly, AAV5 did not transfect cells in the SVZ, confirming it is a valid tool to study the CP-SVZ axis in later experiments (Fig. 4).

With its unique transfection targeted at the CP, we then tested if treatment with AAV5 alone may alter adult neurogenesis in SVZ. By characterizing adult neurogenesis in the SVZ and OB following ICV injection of AAV5 (PBS injections as controls) (Fig. S2A and S2G), we observed that at 30 days after the AAV5 injections, there was no significant alteration in the proliferating NSPCs in the SVZ (Fig. S2B and S2C). Although more Iba-1(+) cells were observed in the CP of AAV5-injected group compared with controls (Fig. S2D and S2E), Iba-1(+) cells were not altered in the SVZ (Fig. S2D and S2F). Similarly, we did not observe any alterations in the density of BrdU(+)/NeuN(+) newborn neurons in GCL or GL (Fig. S2H and S2I) following AAV5 injections. Additionally, as a general indicator of animal wellness, we did not observe any difference in animal’s body weight between groups throughout the transfection (Fig. S2J). Overall, these data supported the AAV5 vector as a “safe” tool for CP-SVZ studies.

AAV5 carrying shRNA targeting Smpd3 were then used to evaluate whether reducing SMPD3 expression in CPECs may suppress sEV release into the CSF. Compared with the controls injected with control vector “AAV5-shRNA(Ctrl)”, in which the shRNA was a non-targeting sequence, injection with AAV5-shRNA(Smpd3) significantly reduced the SMPD3 protein in the CP tissues 4 weeks following the ICV injection (Fig. 3M and 3N). This reduction was further supported by immunohistochemical staining, demonstrating significantly decreased SMPD3 expression in the CPECs (Fig. 3O and 3P). CSF was collected from these mice through cisterna magna puncture, to obtain sEV preparations through differential centrifugation as described previously, and then NTA analysis was conducted. Our data reveal a CPEC-targeted SMPD3 knockdown by AAV5 significantly reduced sEV concentrations in the CSF by 76.7%, compared to the controls (Fig. 3Q). Collectively, we demonstrated SMPD3 was important in maintaining the sEV secretion by the CPECs both *in vitro* and *in vivo*, which justified further investigations to understand how SVZ adult neurogenesis could be modulated by CP through sEVs.

### 3.4. Suppression of CP-derived sEV secretion diminishes SVZ adult neurogenesis and compromises olfactory performance

To address the impact of CP-secreted sEVs on SVZ adult neurogenesis, we quantified the proliferating NSPCs in SVZ one month post ICV injection of AAV5-shRNA(Smpd3) (Fig. 4A). Compared to controls with AAV5-shRNA(Ctrl), a suppressed sEV release by SMPD3 knockdown significantly decreased proliferating BrdU(+) NSPCs in SVZ by 21.3% (*p* < 0.05) (Fig. 4B and 4C). This observation provided direct evidence that CP-secreted sEVs contributed to maintaining the SVZ NSPC activity by sustaining the proliferation dynamics.

SVZ NSPCs ultimately supply newborn neurons to the OB. Thus, we examined the OB newborn neuron density in short term (90 days) and long-term (180 days) post sEV suppression in CP (Fig. 4D, and G). In both experiments, BrdU pulse labeling was performed one month prior to tissue dissection. In the short-term study, suppression of sEV secretion by shRNA(Smpd3) significantly reduced BrdU(+)/NeuN(+) newborn neurons in GCL by 42.2% compared to the controls (Fig. 4E and 4F). Newborn neurons in GL, however, was not significantly affected (Fig. 4E and 4F). Data from the long-term study showed the comparable outcome, i.e., a prolonged suppression of sEV secretion by CP further diminished newborn neurons in OB by 62.0% in GCL but not in the GL (Fig. 4H and 4I). These observations provided additional evidence to support a critical role of CP-derived sEVs in maintaining the SVZ adult neurogenesis.

As the precursors for newborn neurons, DCX(+) immature neurons not only reflect the extent of ongoing neurogenesis (Zhou et al. 2022), but also reveal the synaptic plasticity of the local neuronal network (Marín-Burgin et al. 2012; Schmidt-Hieber et al. 2004). Hence, we quantified DCX(+) cells in the OB 180 days post the CP-targeted transfection. In agreement with our data on diminished BrdU(+)/NeuN(+) cells in the OB, the density of DCX(+) cells was significantly reduced by 81.1% in shRNA(Smpd3)-treated animals (Fig. 4J and 4K). Moreover, the morphology of OB DCX(+) cells appeared to be altered, i.e., in contrast to the typical migratory phenotype such as elongated leading and trailing processes seen in controls, DCX(+) cells in shRNA(Smpd3)-treated animals displayed shortened cellular processes (Fig. 4L). These findings, once again, implicated that altered CP sEVs affected the OB neuronal network composition.

The primary function of the OB is to relay olfactory signals received by the olfactory epithelium to specific receiving areas in brain cortex. To test the olfactory function as affected by CP sEV secretion, we used multiple olfactory testing protocols to assess animals’ olfaction (Kobayakawa et al. 2007; Yang and Crawley 2009), i.e., the buried food test (Fig. 4M), the olfactory preference test (Fig. 4P), and the olfactory avoidance test (Fig. 4S), following short- and long-term CP-targeted sEV alteration. First, when tested at 3 months post injections of AAV5, animals with a reduced CP sEV secretion spent an average of 66.0 ± 19.2 sec in finding the hidden pellet, which was 47.3% longer than animals in the control group (44.8 ± 18.7 sec) (Fig. 4N). The discrepancy was expanded to 180.6% longer in AAV5-shRNA(Smpd3)- treated animals, when the test was conducted at 6 months post injections (Fig. 4O). Data by the pellet-finding test demonstrated that a normal CP sEV secretion was essential to animal’s foraging function.

Second, we conducted the olfactory preference test in male mice using urine collected from female mice by measuring the time for mice to stay with their preferred attractant odorant in a dishabituation session (Fig. 4P) (Kobayakawa et al. 2007). Following the AAV5 injections at 3 months, male mice in the AAV5-shRNA(Smpd3) group investigated the female urine (and staying with it) for 8.8 ± sec, which was 27.2 % less than the controls (12.1 ± 2.2 sec) (Fig. 4Q). This decrease in staying with preferred attractive female urine reoccurred 6 months post the AAV5 injections, but with more extended decrease by 61.0% in AAV5-shRNA(Smpd3)-treated animals than in AAV5-shRNA(Ctrl) controls (Fig. 4R). Thus, the decreased time toward female urine suggested that CP-secreted sEVs were critical in maintaining certain olfaction-based social behaviors.

Third, we tested animals’ avoidance behavior by introducing 2-MB acid (a proven olfactory repellent) to the smaller compartment and thus quantifying their avoidance time in the larger compartment relative to diH_2_O presentation (Fig. 4S). At 3 months post AAV5 injections, control mice in AAV5-shRNA(Ctrl) group were more likely to avoid the 2-MB acid compartment (28.4 ± 9.2 sec relative to water) compared to the AAV5-shRNA(Smpd3) group (17.5 ± 11.0 sec relative to water) (Fig. 4T). Performing the same avoidance testing at 6 months after AAV5 injections, the discrepancy between the 2 groups remained; relative to water, AAV5-shRNA(Smpd3)-treated mice were less likely to avoid the 2-MB acid-present space (22.2 ± 8.7 sec) than were the controls (35.5 ± 6.6 sec) (Fig. 4U). Thus, CP sEV secretion also contributes to danger-escaping scenarios. Taken together, these neurobehavioral experiments support our hypothesis that a diminished newborn neuron supply to the OB, as a consequence of suppressed sEV secretion by the CP, could ultimately deteriorate the olfaction.

Finally, considering the cellular heterogeneity of OB and contributions of various OB cell types to olfaction (Tepe et al. 2018), we characterized the microglia, astrocytes, and oligodendrocyte lineage cells as affected by CP sEV secretion (Fig. S3). By using double-staining of Iba-1 and CD68, we showed that neither pan-microglia (Iba-1^+^) nor activated microglia (Iba-1^+^-CD68^+^) were significantly altered (Fig. S3A and S3B). Similarly, astrocytes in the OB were unaffected by CP-secreted sEVs as the density of S100β^+^-GFAP^+^ cells was not significantly different between the two groups (Fig. S3C and S3D).

Given the importance of oligodendroglia cells in maintaining OB network, we analyzed the myelination status using myelin basic protein (MBP) staining with pan-oligodendroglia marker Olig2 and oligodendrocyte progenitor cell (OPCs) marker NG2 in the OB (Fig. S3E). Quantifying MBP fluorescence intensity in the OB showed that the myelination status stayed unchanged (Fig. S3F). The Olig2(+) cell density was also unaltered by suppressed CP sEV secretion (Fig. S3G). Moreover, we did not observe a significant change in NG2(+) OPCs in the OB (Fig. S3H); yet the OPC distribution pattern seemed to change toward more OPCs in the core of OB in the AAV5-shRNA(Smdp3) group in contrast to a GCL-dominant presence in controls. Overall, a lack of evident alterations of glial cell populations in the OB supports our hypothesis that the diminished SVZ adult neurogenesis by a suppressed CP sEV secretion drives the deterioration of olfactory function.

## 4. Discussion

The data presented in this assessment provide direct evidence to support the existence of a CP-SVZ regulatory (CSR) axis, in which the CP regulates the adult neurogenesis in SVZ through its production and secretion of sEVs. Further, the proposed CSR axis is significantly involved in maintaining normal olfactory function. The concept of the CSR axis is supported by several lines of evidence in this evaluation. First, sEVs were identified to be the critical fraction in the CP secretome stimulating SVZ neurosphere growth. Additionally, impaired newborn neuron supply to the OB following ICV infusion of sEVs collected from the CP of Mn-exposed animals further supports this concept. Second, by revealing the critical role of SMPD3 in determining sEV release from CPECs *in vitro* and using AAV5 for CP-targeted SMPD3 knockdown *in vivo*, we observed the secretion of sEVs by the CP determined the sEV concentration in the CSF. Third, selective suppression of CP sEV secretion by SMPD3 knockdown progressively diminished adult neurogenesis in the SVZ-RMS-OB system. Finally, neurobehavioral tests demonstrated alteration in the CP secretion of sEVs ultimately led to impaired olfactory function. Collectively, our findings highlight the importance of the CP, through sEV secretion, in modulating the adult SVZ neurogenesis process, thereby influencing the OB/olfaction system.

The CP-associated secretome contains health- and growth-promoting factors that are critical to the wellbeing of developing and adult brains. Literature indicates substances released by the young CP are capable of “rejuvenating” old brains by upregulating brain plasticity (Iram et al. 2022; Silva-Vargas et al. 2016). Our data are in agreement with these reports, and further demonstrate the sEVs secreted by the CP may mediate these beneficial effects. Interestingly, many “nourishing” molecules in the CP such as Klotho (Grange et al. 2020; Zhu et al. 2018), microRNA-204 (Lepko et al. 2019), SOD3 (Abdelsaid et al. 2022; Jang et al. 2022), sonic hedgehog protein (SHH) (Lun et al. 2015; Vyas et al. 2014) likely exert their biological functions using sEVs as the dominant carrier. The CP’s rich blood supply makes it sensitive to systemic biological or chemical changes in the body; hence it is possible the CP produces and utilizes sEVs as a primary mechanism to communicate with cells within the SVZ. This CP-SVZ signaling is promoted via the pinwheel architecture of the SVZ on the ventricular surface which can directly contact with the CSF (Mirzadeh et al. 2008). The “antenna” or a subcellular structure known as the primary cilium (Tong et al. 2014) in SVZ NSPCs facilitate sEV uptake for subsequent signal reception/transduction (Corbeil et al. 2020; Hogan et al. 2009). The exact interactions between CP-derived sEVs and cellular components of the SVZ remains unknown and requires extensive future investigation. However, the CP’s robust sEV secretion and the SVZ’s high sensitivity to changes in the CSF appear to collectively underpin the physiological existence of the CSR axis. Understandably, other mechanisms must exist; for example, orthodenticle homeobox 2 (OTX2), a critical neuronal lineage transcription factor enriched in the CP, likely enters the CSF directly as a soluble protein (Bernard et al. 2016; Planques et al. 2019). Copper (Cu) in the CSF, regulated by the transporters in the CP, can also modulate adult neurogenesis in SVZ not necessarily via CP-derived sEVs (Liu et al., 2022).

Clinical implications of the CSR axis remain to be elucidated. Significant, however, is the linkage of the proposed CSR axis to olfaction. Olfactory disorders represent a highly prevalent early-onset symptoms in several neurodegenerative disorders, particularly in Alzheimer’s disease (Murphy 2019; Pacyna et al. 2022; Tian et al. 2022) and Parkinson’s diseases (PD) (both idiopathic and Mn-induced) (Doty 2012; Lucchini et al. 2009). Disturbed olfaction is also one of the prominent leading symptoms associated with post-COVID conditions (Boscolo-Rizzo et al. 2022; Mendes Paranhos et al. 2022). Interestingly, severely disrupted function of the CP in humans has been associated with pathological progressions in AD, frontotemporal dementia, Huntington’s disease, and severe COVID19 (Kant et al. 2018; Stopa et al. 2018; Tadayon et al. 2020; Yang et al. 2021). Thus, our proposed CSR axis may provide a useful explanation to connect the CP’s sensitivity to changes in the circulation’s chemical environment to olfactory dysfunction. It is possible the CP may directly communicate with the OB through secreted CSF, which exchanges freely with the interstitial fluid between neurons and glial cells in brain structures (Zheng and Chodobski 2005). Alternatively, the CP may influence olfaction through the immediately adjacent SVZ NSPCs as proposed in this study. Noticeably, there was a debate in literature regarding the importance of the SVZ adult neurogenesis in human olfaction (Curtis et al. 2007; Sanai et al. 2007). Our findings, however, provide compelling evidence to support the sEV-dependent CSR axis in modulating the olfactory sensation. We anticipate “single-cell” techniques, i.e., single-cell/single-nucleus RNA-sequencing, may further confirm ongoing neurogenesis in adult human olfactory epithelium (Durante et al. 2020) and uncover interspecies differences in adult neurogenesis (Zhou et al. 2022, 2023).

The concept of the CSR axis has a significant implication in toxicology. The ability of the CP to effectively sequester toxicants, owing to its interactions with blood-borne extrinsic materials, renders the tissue vulnerable to chemical-induced injury not solely on its structure but also on its function (Fu et al. 2014; Ingersoll 1995; Zheng 2001; Zheng et al. 1991, 1999). The data presented in this study supports exposure to Mn leads to extensive accumulation of Mn in the CP, which in turn alters the signals carried by sEVs from the CP. At present, the evidence to support toxicant-induced dysfunction of sEV secretion by the CP with ensuing impaired olfaction remains scarce. Nevertheless, the CSR axis may become a new target for toxicological understanding of chemical-induced brain injury. In addition, the CSR axis may provide new mechanisms and new targets for pharmacological intervention to address multiple neurological diseases. Multiple clinical trials have shown a promising therapeutic outcome of sEV-based interventions (Besse et al. 2016; Nassar et al. 2016; Perocheau et al. 2021; Shapira et al. 2022; Zhu et al. 2022). Factors secreted by the youthful CP appears to improve brain function (Baruch et al. 2014; Iram et al. 2022; Silva-Vargas et al. 2016); hence, youthful factors secreted by the CP derived from induced-pluripotent stem cells (hiPSCs) could be harnessed for experimental trials (Jacob et al. 2020; Pellegrini et al. 2020; Sakaguchi et al. 2015).

In conclusion, the data presented in this study provide first-hand evidence to support the existence of the Choroid plexus-Subventricular zone Regulatory (CSR) axis, which, through the yet-to-be-identified molecules carried by CP-secreted sEVs, regulates adult neurogenesis in the SVZ and ultimately modulates the olfactory function. Mn-caused anosmia/hyposmia may be due at least partly to its adverse effects on the CSR axis. Further defining of the CSR axis will have significant implications in neurobiological, neurodevelopmental, neuropharmacological and neurotoxicological research and application.

## Limitations of the study

This report lacks cargo content analyses of CP-secreted sEVs. Analysis of CP-secreted sEV cargo shall further elucidate the mechanism by which Mn-sEVs differentially modulate neurogenesis to the OB, and in the meantime, help identify SMDP3-dependent secreted cargo. In addition, this study used male animals only to assess the olfaction as mediated by CP-secreted sEVs, taking into account the susceptibility of the males to olfactory disorders. Future research should include the neurobehavioral characterizations in less susceptible female animals. Ongoing and future studies will further elucidate this mechanism by examining specific sEV cargo components influencing neurogenesis, sEV targeting systems, and disease- as well as sex-mediated variations to establish clinical targets. Overall, our study represents foundational knowledge supporting the CSR axis and the contribution of CP-derived sEVs.

## Acknowledgement

This study is supported by NIH R01 ES027078. The authors thank Richard van Rijn at Purdue University for the stereotaxic device, Christopher Gilpin and Laurie Mueller at Life Science Microscopy Facility at Purdue University for their technical guidance for electron microscopic analysis, Jason R. Cannon at Purdue University for the microtome uses, and Ariel Di Nardo and Alain Prochiantz at Collège de France for their technical suggestions for the selection of AAV5. Illustrations in this report were created with the templates in BioRender.com.

## Author contributions

Conceptualization, W.Z., L.L.; Methodology, L.L., W.Z., J.S.; Investigation, L.L., W.Z.; Analysis, L.L., W.Z., J.S.; Original Writing, L.L., W.Z.; Review & Editing, L.L., W.Z., J.S.; Supervision, W.Z.; Funding Acquisition, W.Z.

## Supplemental Figures and Legends

**Figure S1.**
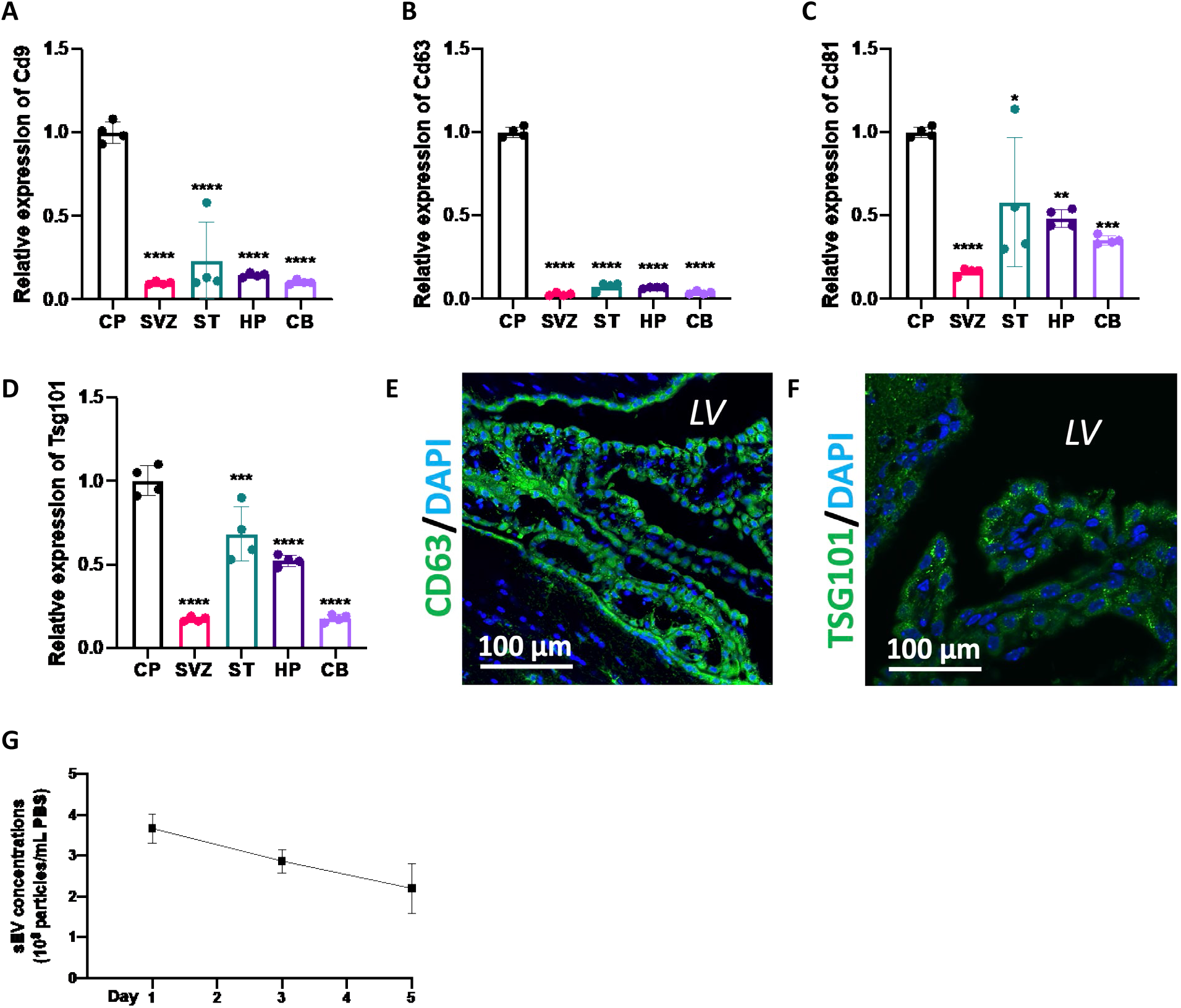
CP expresses high level of sEV markers. (**A-D**). qPCR analyses for relative expressions of sEV markers across brain regions. Data of Cd9, Cd63, Cd81, and Tsg101 were provided in (A), (B), (C), and (D), respectively (n = 4). ST: striatum; HP: hippocampus; CB: cerebellum. (**E**). IHC staining of CD63 in the CP. (**F**). IHC staining of TSG101 in the CP. (**G**). Effects of storage duration under 37°C on concentration of sEVs (n = 3).

**Figure S2.**
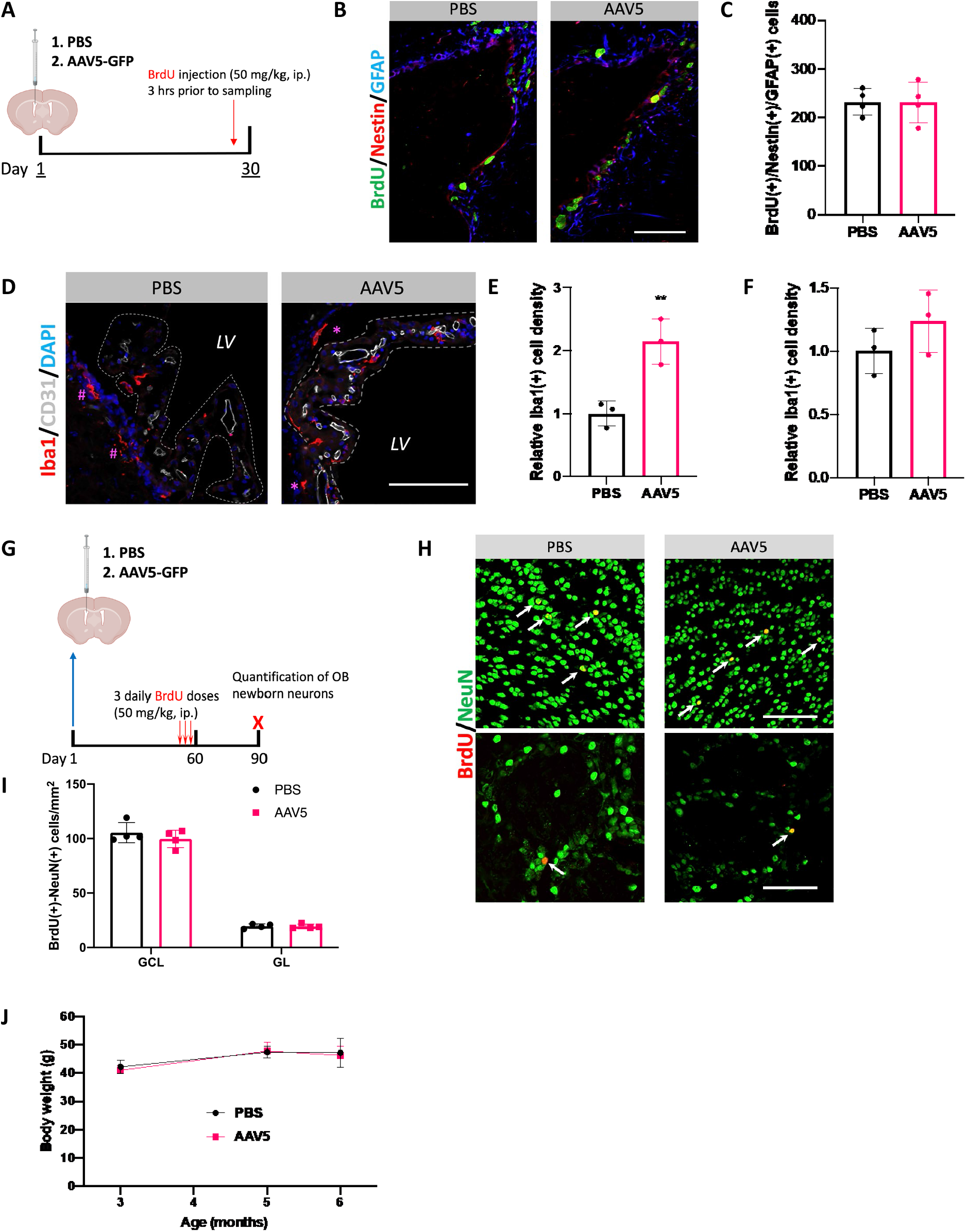
AAV5 does not evidently impact SVZ adult neurogenesis. (**A**). Experimental timeline assessing the effects of AAV5 ICV injection on proliferation of SVZ NSCs. (**B** and **C**). Representative images and quantification of proliferating SVZ NSCs 30 days post ICV injection of AAV5. Of note, mice used in this experiment were 5-month-old. n =4; scale bar = 100 µm. (**D**-**F**). Representative images and quantification of Iba1(+) cells in the CP and SVZ 30 days post ICV injection of AAV5. Relative Iba1(+) cell density in the CP and SVZ were provided in (E) and (F), representatively. n =3; scale bar = 50 µm. (**G**). Experimental timeline assessing the effects of AAV5 ICV injection on SVZ adult neurogenesis to the OB. (**H** and **I**). Representative images and quantification of BrdU(+)/NeuN(+) newborn neurons in the OB. In (H), the upper panel shows newborn neurons in the GCL, and the lower represents newborn neurons in the GL. n =4; scale bar = 50 µm. (**J**). Body weight monitor post ICV injections (n = 5).

**Figure S3.**
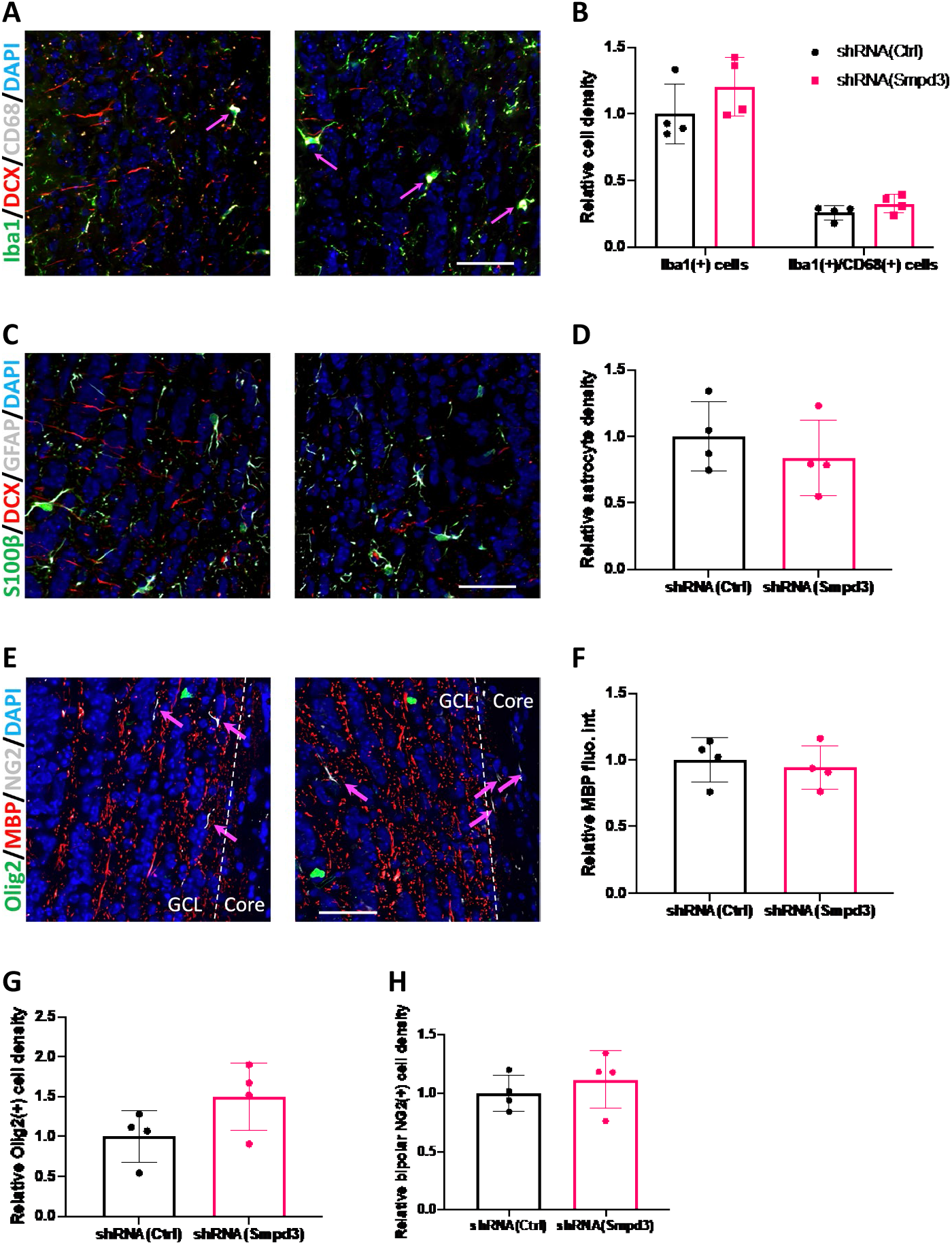
Glial population in the OB 6 months post CP-targeted SMPD3 knockdown. (**A** and **B**). Representative images and quantification of Iba1(+) microglial cells and the CD68(+) activated phenotypes in the OB. n =4; scale bar = 50 µm. Arrows in pink mark enlarged Iba1(+)/CD68(+) cells. (**C** and **D**). Representative images and quantification of GFAP(+)/S100β astrocytes in the OB. n =4; scale bar = 50 µm. (**E**). Representative images of oligodendrocyte lineage cells in the OB. Scale bar = 50 µm. Arrows in pink mark bipolar NG2(+) cells. (**F-H**). Quantification of MBP fluorescent intensity, Olig2(+) cell density, and NG2(+) cell density in the OB. n =4.

